# A neural framework for spontaneous development of orientation selectivity in the primary visual cortex

**DOI:** 10.1101/2022.09.06.505990

**Authors:** Catherine E. Davey, Errol K.J. Lloyd, Levin Kuhlmann, Anthony N. Burkitt, Trichur R. Vidyasagar

## Abstract

The mechanism or microcircuitry behind orientation selectivity in primary visual cortex (V1), and the means by which it develops without supervision or visual input, both remain unresolved questions. Work on the developmental question has assumed the prevalent spatial convergence model of orientation selectivity as the target mechanism. Encouraged by growing evidence challenging both the completeness of this model and its developmental viability, we investigated an alternative scheme. Accordingly, we demonstrate computationally how a scheme in which orientation selectivity originates from the orientation biases already in the retina and lateral geniculate nucelus (LGN) can answer both the mechanistic and developmental questions. In this scheme, the divergence of outputs from the retina allows retinal spontaneous activity to create correlations within the LGN. These correlations in turn allow a Hebbian plasticity mechanism to strengthen those LGN inputs to V1 which carry similar orientation biases and thus provide an orientation tuned excitatory input.

## 1 Introduction

The hallmark of most neurones in the primary visual cortex (area V1) is that they respond selectively to stimuli of a particular orientation^1,2^. The neural basis of this selectivity is still under much debate^3–5^. The earliest proposal^1,2^ was that a primary visual cortex (V1) neuron receives multiple inputs from the lateral geniculate nucleus (LGN) with receptive fields (RFs) that are spatially spread along a particular line or axis, such that maximal input is generated by stimuli with edges aligned along the same axis. The core of this scheme is the convergence of receptive fields that are spatially aligned, called ‘spatial convergence’. This scheme, however, has issues with explaining all the behaviours of V1 neurones and so is unlikely to be complete or accurate.

One long-standing alternative scheme is based upon the fact that sub-cortical cells in the retina ^6,7^, and resultantly the LGN^6,8,9^, are also orientation selective, albeit to a lesser degree compared to V1 cells. This ‘sub-cortical biases scheme’ proposes that this lesser selectivity (or ‘bias’) in LGN is the source of orientation selectivity in V1, whose chief function is to inherit, filter and amplify already selective inputs^10^. The scheme enjoys both empirical^11–14^ and computational^15^ support and an explanatory capacity that spatial convergence lacks^3,15^. Moreover, recent work shows that the degree of spatial dispersion amongst the LGN cells providing inputs to V1 orientation columns is minimal^16^. This finding is a serious issue for spatial convergence proposals and further raises the fundamental importance that sub-cortical biases may have for V1 function.

Our interest here is how the scheme for orientation selectivity, whichever scheme it is in reality, develops from the embryonic state, a question that has also remained elusive. As V1 achieves orientation selectivity before the eyes open,^17–21^ the development process must rely on some spontaneous and unsupervised mechanism. The most likely origin of a mechanism is the spontaneous activity that cells in the visual pathway produce throughout development. Correlations in this activity may cause the strengths of V1 to LGN synapses to be modified through Hebbian-like plasticity, while patterns in these correlations can then provide the seed for structure in the V1 to LGN circuitry. The challenge, then, for both of the above schemes of orientation selectivity, is to provide an accompanying proposal for how a particular pattern in the spontaneous activity occurs and how it can lead to the requisite circuitry.

Much of the previous computational work on the developmental question has focused on the spatial convergence scheme. Another line of computational modelling of orientation selectivity is one that has focused on the convergence of ON and OFF inputs with offset receptive fields^22–24^ However, in order to achieve the required arrangement of receptive fields, it was necessary in both cases to presume certain patterns of correlation or an anatomical arrangement that have not found empirical support^3,25–27^.

The sub-cortical biases scheme, on the other hand, simplifies the question. For this scheme to develop, the pattern required in the spontaneous activity of the LGN inputs is that those that are biased for the same or similar orientations are more correlated with each other than those biased for dissimilar orientations. Aiding in the creation of such correlation is the divergence of the retinogeniculocortical pathway, which creates a number of either LGN cells (in the case of the cat) or layer 4 cells (in the case of the macaque) transmitting signals from a single retinal ganglion cell^28–37^. Through plasticity, the set of inputs that are then strengthened, which would be those that are more correlated, would differ only slightly or not at all in their orientation biases and so provide a net input that is itself orientation selective. This similarity of selectivity amongst the inputs to individual V1 cells is necessary for the sub-cortical biases scheme, as otherwise the input biases would cancel and there would be no selectivity in the input to V1 cells that they could inherit.

Remarkably, such a developmental process - the grouping of similar inputs - is common in V1, having been demonstrated for a number of response properties. More importantly, the structure of the visual pathway appears to be distinctly conducive to such grouping through a spontaneous activity driven plasticity process. This general structure involves there being multiple cells in the cat LGN for each retinal cell, and each LGN cell being driven largely by a single input retinal cell. That is, the retinal input to the LGN is not only the dominant input, but it is also divergent. Thus, LGN cells are near clones of their ‘parent’ retinal cells in terms of response properties such as orientation selectivity, and those that share retinal input are highly correlated with each other. In the macaque, this divergence happens not so much at the retinogeniculate projection, but rather at the LGN-V1 projection^30^.

Accordingly, compared to the challenges faced by the spatial convergence scheme in answering the developmental problem, our sub-cortical biases scheme may address it with relative ease. It has direct precedents, requires no unfounded presumptions, relies on well-established phenomena and already has supporting evidence^6,8,9,11,13,29,30,38^. Accordingly, here we evaluate and model the viability with which retinal spontaneous activity and Hebbian plasticity may lead to V1 orientation selectivity. We show that the divergence of signals from a single retinal ganglion cell generates a robust self-organised grouping of similarly tuned inputs to a set of V1 cells, which, with simple intra-cortical inhibition, leads to realistic cortical orientation tuning.

## 2 Results

We propose that mild orientation bias in the retinal receptive fields precipitates orientation selectivity in the visual cortex, with inhibitory interneurons being crucial to sharpening the functional orientation bias that V1 receives from the thalamic input. In accordance with the mild orientation bias found in retinal ganglion cells (RGC) cells^6,8,9,39^, and the divergent connections found from the retina to the LGN^35,37,40,41^, we simulated a layer of mildly oriented RGC cells, each connected to several LGN cells (see Fig. 1). Each LGN cell received eight RGC inputs with a Gaussian synaptic connectivity distribution centred on the LGN cell, having a radius of 1.5 RGC cells ^31^. Thus an LGN cell tended to inherit the mild orientation selectivity of the closest RGC cell, which was expected to have the largest number of synaptic inputs.

**Figure 1:**
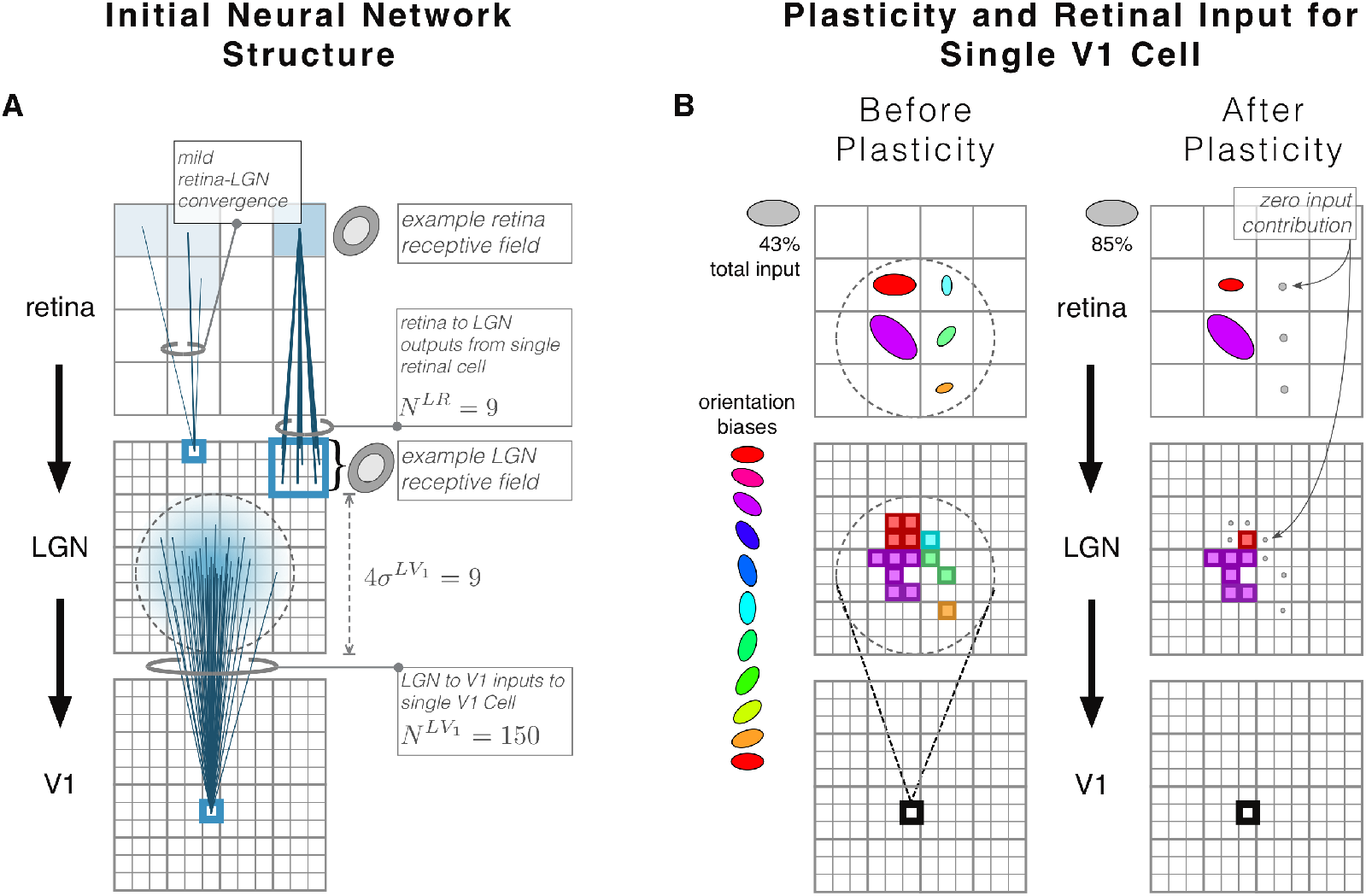
Illustrative depiction of the structure of our three layered feed forward network (a), as well as a depiction of the plasticity process that we propose and demonstrate (b). **(a):** Initial state of the network prior to the simulation of plasticity and development. Three layers of neurons, from top to bottom, representing RGCs. LGN and V1 respectively. Each square represents a cell, and cells are distributed evenly across the layers. Essential aspects of the network are depicted with relevant variables, for illustration purposes only. For an accurate and complete account of the system, see Table. 1 and Section 4. Outputs from the retina to the LGN diverge, governed by the parameters *N^RL^* and *σ^RL^*. *N^RL^*=8, to designate a single RGC cell having 8 synaptic connections to the LGN layer. The synaptic connections are Gaussian distributed with radius *σ^RL^* = 1.5. Given that each LGN cell receives an average of 8 synaptic inputs within a small radius, each LGN cell is dominated by the nearest RGC cell, and thus obtains a receptive field with an orientation bias similar to that of their closest RGC. Additionally. neighbouring LGN cells sharing RGC input tend to shared an orientation bias, since the LGN cell density is greater than V1, so that LGN cells often had inputs dominated by the same RGC cell. While connections from RGC to LGN are divergent, those from LGN to V1 are convergent, with the number of synaptic connections received by each V1 cell denoted by *N^LV_1_^* = 150. Again, the synaptic distribution is assumed to be Gaussian, with a standard deviation *σ^LV_1_^* = 5, depicted by the dashed circle. A single LGN cell may provide multiple inputs, or synapses, to a single V1 cell. **(b):** Proposed plasticity process depicted for a single V1 cell, emphasising the relative contributions of retinal cells to V1, via the LGN layer, and the consequent RGC effect on the plasticity process. Layer structure is identical to that in (a), and the orientation biases of retinal cells contributing to the example V1 cells are depicted by both color and ellipse orientation. Due to the randomness of the initial synaptic connections between layers, an RGC cell may have multiple synaptic inputs to a single LGN cell, and may also be connected to a V1 cell via multiple LGN cells, Consequently, the contribution of each RGC to a single V1 cell varies, and is depicted by variation in ellipse size (see also reference size at top left of retinal layer). The LGN cells via which retinal inputs are relayed to V1 are colored according to their dominant retinal input. Not shown here is that LGN cells can vary in the number of synapses they provide to a given V1 cell. The purple RGC initially provides the most input, so that it dominates activity in the V1 cell. Consequently. there is stronger correlation between the V1 cell and the LGN cells relaying the purple RGC activity to V1. Plasticity therefore favours the LGN to V1 synapses that relay the purple retinal and LGN input while other LGN inputs (blue, green and orange) are extinguished by plasticity (depicted by grey circles, After plasticity, the input to the V1 cell would bear an orientation bias derived from that of the purple retinal cell.

The distribution of orientation preferences in the retina was adopted from experimental data^39^, drawn from 10 unique orientations. A cortical V1 neuron received excitatory inputs from multiple LGN cells, with a random Gaussian connectivity distribution centred on the V1 cell.

To examine the mechanism behind the sharpened orientation selectivity found in V1 we simulated a network with only the feed-forward connectivity discussed and a second network in which there were additional lateral inhibitory interneurons in the V1 layer. Inhibitory connections to a V1 neuron were random with a Gaussian distribution larger in spatial extent than the excitatory connections from the LGN. The inhibitory neurons, being located in the V1 layer received the same input as the excitatory neurons in V1, namely the LGN cells. We refer to the networks as the feed-forward and lateral-inhibitory networks respectively.

In addition to considering the mechanism behind orientation selectivity in V1 we also considered its development via the well-established principles of Hebbian plasticity^22,42–44^. Since orientation selectivity is known to be established before the onset of structured environmental input, we simulated its development in the presence of spontaneous, Poisson noise in the retina. The strength of synaptic connections between the LGN and V1 were allowed to evolve according to a Hebbian learning equation (see Eqs. (4) and (5)), until all weights had either decayed to 0 or reached the upper bound.

On completion of the development phase we evaluated the orientation and spatial frequency tuning of the V1 cells of each network. We also examined the receptive fields of the V1 cells for evidence of spatial dispersion along an axis, as required for the spatial convergence scheme of orientation selectivity ^1^. Additionally, we evaluated each of these measures after 30% of the simulation had completed, where the length of the simulation was determined by the time taken for the synaptic weights to converge to stable values, thus being able to compare three stages during development: ‘at birth’, ‘during plasticity’ and at ‘maturity’. The value of 30% was chosen to approximately match the results of Kiley and Usrey^45^, in which the RF size of cells in response to white noise were recorded at 7 days after birth and in the adult cat. Since we do not have any data about retinal, LGN or cortical RFs at birth, we started the ‘at birth’ stage assuming a certain degree of convergence from retina to single LGN cells and then let the simulation run for a while to match, approximately, the earliest detailed data for receptive fields available at 1 week (before eye opening in the cat) from the work of Kiley and Usrey^45^. According to their data, the RFs at 1 week are about 4 times the size the adult RFs (from their Fig. 3 and the data at less than 5 deg eccentricity). However, this ratio requires an important correction, since the relevant measure is the real centre size, but the effective RF sizes of both LGN and visual cortex measured by Kiley and Usrey^45^ in the adult cat are constrained by inhibition unlike in the newborn, a point that was not considered by the authors in their comparison of RF centre diameters between 1 week old and adult cats. In the kitten, experiments using bicuculline iontophoresis to block gamma-aminobutyric acid (GABA) receptors show that inhibition in the LGN does not start becoming functional till 45 days after birth and reaches adult strength only by 100 days^46^. Thus the actual excitatory discharge region of adult LGN cells are likely to be larger than those in the recordings of Kiley and Usrey^45^. Thus we aimed for a ratio of two (instead of 4) between the 1 week and adult LGN and V1 RF sizes. This may seem somewhat arbitrary, but it accounts for the likely underestimation of adult centre diameters by Kiley and Usrey^45^. LGN RF sizes measured by Daniels et al.^47^ at 1 week and 4-5 week old kittens also show the gradual reduction, which is probably due at least partly to the maturation of inhibition. Inhibition in V1 is also poorly developed in the newborn and is shown to mature only by four weeks after birth ^48^. Using the above results as a guide, we generated RFs of V1 cells at 30% of the total simulation duration, because V1 cells response to white noise generated RFs with approximately twice the size.

**Figure 2:**
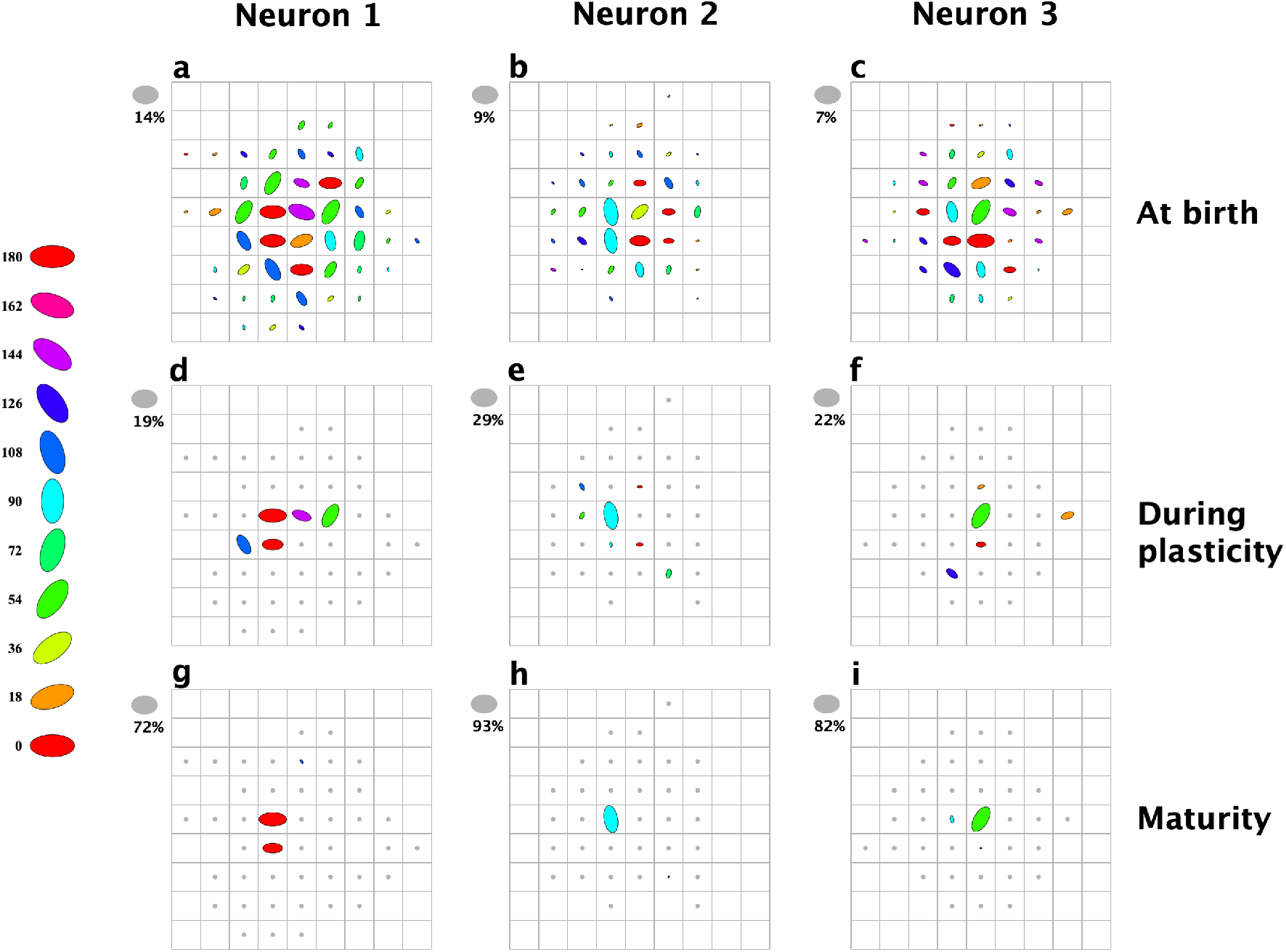
Feed-forward network: Shows the RGC inputs to three exemplar V1 neurons, mediated by the LGN. The RGCs are arranged uniformly across the laminar. The orientations of the retina (retina) cells are indicated by an ellipse - a cartoon depiction of the DOG kernel defined in Eq. (7) - rotated to the appropriate orientation. Each orientation is also allocated a unique colour, as indicated by the legend. Each retina cell can have multiple connections to a LGN cell, and a LGN cell may have multiple connections to a particular V1 cell, essentially inputting multiple copies of the retina cell’s activity to the V1 cell. The percentage of V1 presynaptic connections from each retina cell to a V1 neuron is depicted by the size of the ellipse, and the maximum percentage for each V1 cell is shown with the grey ellipse to the left. Parameters governing the development of synapses between the retina and LGN, Eq. (4a), were chosen such that all synapses reached the upper bound. Parameters for the development of synapses between the LGN and V1, Eq. (4b), were chosen such that ⅕ of the initial connections in to a V1 cell were expected to survive after the cell had reached maturity. The expected number of surviving synapses was determined by the homeostatic equilibrium, described in Eq. (6), for which the parameters were set to 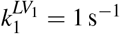 and 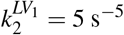. **(a-c):** Presynaptic connections from the RGC layer to three V1 neurons, mediated by the LGN, before development. **(d-f):** The same three neurons during plasticity, but prior to convergence of weights at maturity. Figures show the number of presynaptic connections from the LGN after 30% of the simulation has run. **(g-i):** The same three neurons after development. Figures show the number of presynaptic connections from the LGN that survived plasticity, reaching the upper bound of 1.

**Figure 3:**
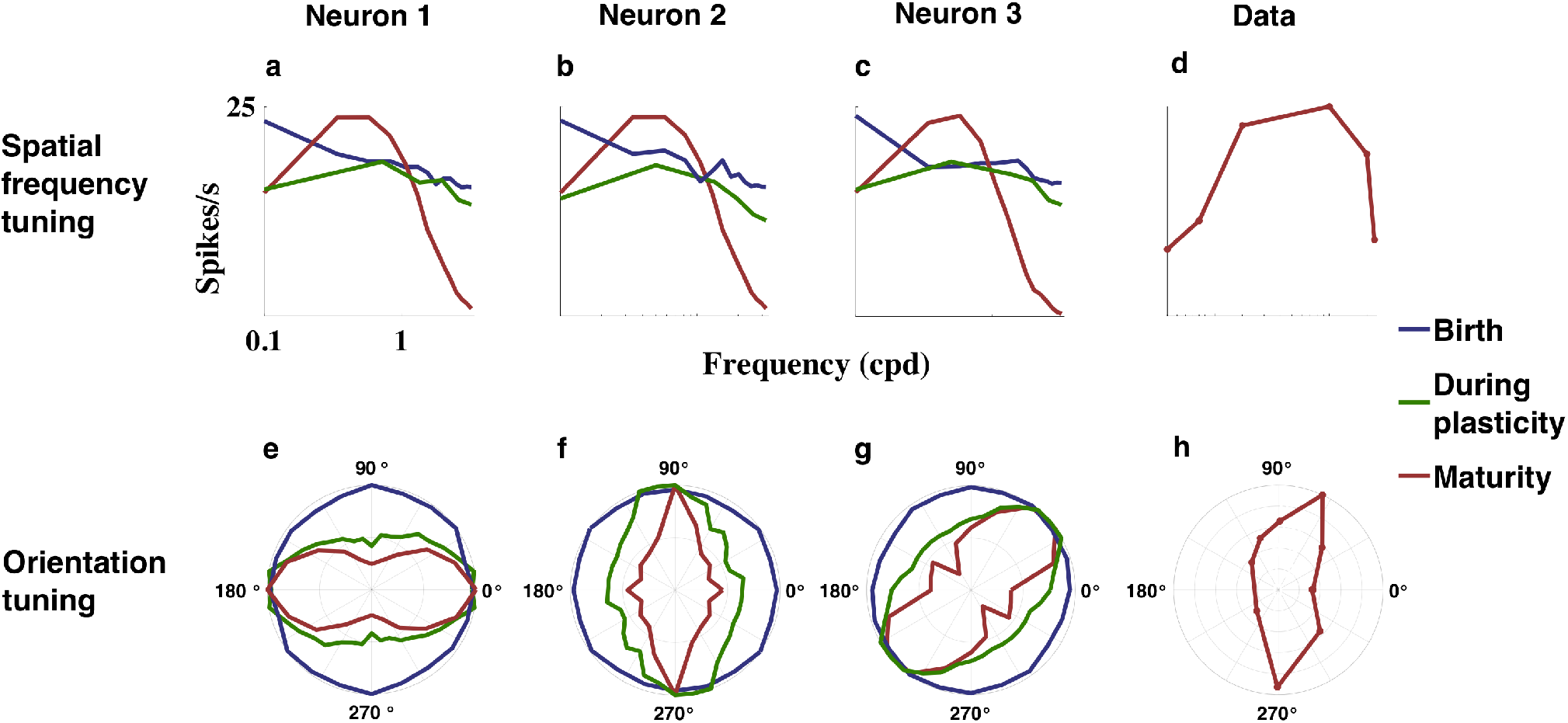
Tuning curves of V1 cells from the feed-forward network. The network was stimulated with moving sinusoidal gratings, detailed in Eq. (8), to evaluate functional orientation and spatial frequency selectivities of V1. A V1 cell’s maximum firing rate was recorded in response to each stimulus orientation and spatial frequency combination. Tuning curves are shown for three examplar cells in V1 of the feed-forward network prior to learning at birth (blue), during the learning process (green), and at maturity after weights have converged (red). **(a-c):** Spatial frequency response of three feed-forward network V1 cells. **(d):** Spatial frequency response from a cat V1 cell^49^. Responses were measured following suppression of inhibitory synaptic weights using bicuculline, a GABA_A_ antagonist. **(e-g):** Normalised orientation tuning curves of the three feed-forward V1 neurons. **(h):** Normalised orientation tuning curve of a cat V1 cell after suppression of inhibitory synaptic weights using bicuculline, a GABA_A_ antagonist.

The networks were driven by unstructured Poisson noise from the retina, and both LGN and V1 cells were modelled as rate-based Poisson neurons according to Eqs. (2) and (3), respectively. Connectivity between layers in each of the networks was randomly initialised according to a Gaussian connectivity kernel of specified radius (see σ values in Table. 1). Synaptic weights were randomly initialised using a uniform distribution between 0.1 and 0.3 (see Table. 1 for details). While inhibitory connections were assumed constant, excitatory connections were allowed to evolve during development according to the weight update dynamics described in Eqs. (4) and (5). Since the plasticity equations are unstable^42^, synapses either decayed to the lower bound of 0, or grew to the upper bound of 1. As in Linsker^42^, excitatory synapses between the first two layers - between the retina and the LGN - are all assumed to evolve to the upper bound of 1. Connections to deeper layers were evolved only after all synapses in earlier layers had reached a bound.

**Table 1:**
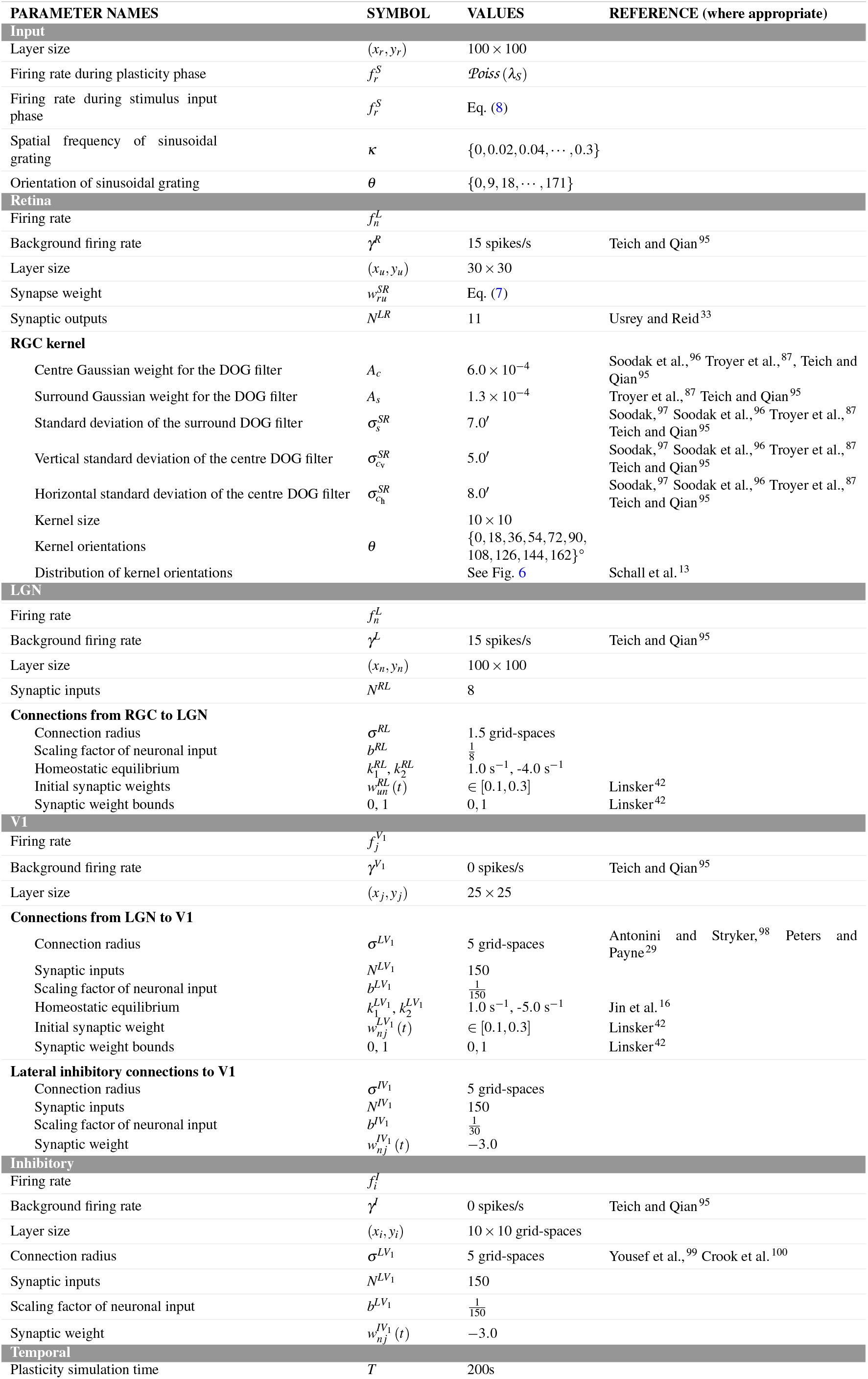
Network parameter names, values, and references.

Note that throughout this section, figures depicting results for individual neurons show the same three exemplar neurons from the V1 layer of each network.

### 2.1 Feed-forward network

#### 2.1.1 Development

RGC inputs to three exemplar V1 cells from the feed-forward network are depicted in Fig. 2, shown both before, during and after development. V1 cells receive RGC activity via the LGN. As each LGN cell has eight RGC inputs from within a small Gaussian connectivity radius of 1.5 cells, it inherits the mild orientation bias of the nearest RGC cell, which is expected to have the most synaptic input.

The bias is modelled by a difference of Gaussian (DOG) filter, Eq. (7), which is illustrated graphically using an ellipse, rotated to depict the orientation of the RGC cell. Additionally, each of the 10 possible orientation biases was allocated a unique colour, and the RGC cells coloured according to their orientation to show the range of input orientations to the V1 cell. A V1 neuron may receive multiple copies of an RGCs’s activity if it has connections from LGN cells with the same pre-synapsing RGC. The number of copies of an RGC cell that are received by the V1 cell is depicted by the width of the ellipse.

Fig. 2 shows the number of distinct retinal inputs to a single V1 cell that survive after plasticity and Fig. 3 shows the orientation tuning that emerges after plasticity. Any orientation bias evident in the V1 layer after development does not result from the spatial convergence of LGN receptive fields, since the preferred orientation of these neurons, as evident in Fig. 3, have little resemblance to the orientation that any apparent spatial convergence, as seen in Fig. 2, may suggest. Rather, LGN cells of the same, or fairly similar orientation preference are seen to survive development and hence come to dominate input to a V1 cell. This is achieved by a substantial reduction in the number of input orientations during development. LGN cells with the same presynaptic RGC cell will have correlated input. The Gaussian synaptic connectivity distribution ensures that LGN cells more proximate to the V1 neuron will have more RGC cells in common, increasing covariance with presynaptic neurons, and conferring on such cells a competitive advantage during development. Hence LGN cells with the most RGC cells in common will typically survive first. Given that the number of mature connections depends on the homeostatic constants, if the number of LGN connections from the dominant RGC cell are fewer than the expected number of mature connections (see Section 4.2), then LGN cells from the next most dominant RGC cell will typically survive next, not withstanding random fluctuations, and so on. Consequently, the number of input orientations will depend on the homeostatic constants, *k_1_* and *k_2_*, which determine the number of synapses expected to survive, and the synaptic connection radius, which determines the expected number of connections from each LGN cell to the V1 cell. Distinct RGC cells with the same orientation will only gain a competitive advantage from each other if their receptive fields overlap so their neural activity is correlated.

#### 2.1.2 Tuning of V1 cells

After establishing a lack of spatially induced orientation bias, we evaluated the V1 cells in both networks for evidence of functional orientation selectivity. Orientation and spatial frequency tuning curves were generated by inputting moving sinusoidal gratings of specified spatial frequency, *κ*, and orientation, *θ* (see Table. 1 for possible values). The tuning curves generated from the model networks were qualitatively compared to the tuning curves of real V1 cells obtained experimentally from the cat visual cortex^49^. In order to meaningfully compare the response of the feed-forward network containing only excitatory synapses, we used data obtained after iontophoretic application of bicuculline, a GABA antagonist that suppresses inhibitory weights by a factor of 0.1 ^15,49,50^ (Fig. 3d).

Spatial frequency and orientation tuning curves for V1 neurons from the feed-forward network are displayed in Fig. 3. Note that the V1 neurons in the figure are the same neurons as shown in Fig. 2. The tuning curves were generated from each cell’s neural activity in response to moving sinusoidal gratings, described in Eq. (8). Fig. 3a-c show the spatial frequency responses of the simulation neurons from the feed-forward network to their preferred orientation both before, during and after development. It can be seen that before plasticity has selectively culled synapses, the large range of input orientations effectively combine to create an unbiased receptive field. Consequently, the lower spatial frequencies evoke a higher response rate. After development, less dominant orientations have been culled, leaving only a few RGC cells connected to the V1 cell, and therefore only a few input orientations. Consequently, there is no longer a spatial averaging over many input orientations, and the peak of the spatial frequency curve after learning corresponds to the spatial frequency of the RGC DOG kernel. Included in panel D is the tuning curve of a real V1 cell.

Fig. 3e-g, show the orientation tuning curves for the three neurons in the V1 layer of the feed-forward network, before, during and after development. Before plasticity, when there were a substantial number of input orientation biases originating from many different RGC cells, the V1 cells responded maximally to all grating orientations. After development there was typically a dominant orientation derived from many connections from a single RGC cell. Consequently, the orientation tuning curves post plasticity had similar circular variances (CVs) to that exhibited in the retina, shown in Section 4.3. Thus, Fig. 3 shows that orientation selectivity in the cortex can be acquired from orientation bias in the retina.

### 2.2 Lateral-inhibitory network

#### 2.2.1 Development

The lateral-inhibitory network that we simulated incorporates lateral inhibitory interneurons in the V1 layer. These connections were not plastic, and synaptic weights were initialised to −3 (see Table. 1). RGC inputs to three exemplar V1 cells from the lateral-inhibitory network are depicted in Fig. 4, shown before, during and after development. See Section 2.1.1 for an explanation of the corresponding figure for the feed-forward network. The excitatory connections from the LGN to the inhibitory interneurons within V1 were modelled as having the same statistics as the excitatory connections from LGN to V1 in the feed-forward pathway. The inhibitory connections to V1 were also Gaussian, having a larger radius than the excitatory connections to V1.

**Figure 4:**
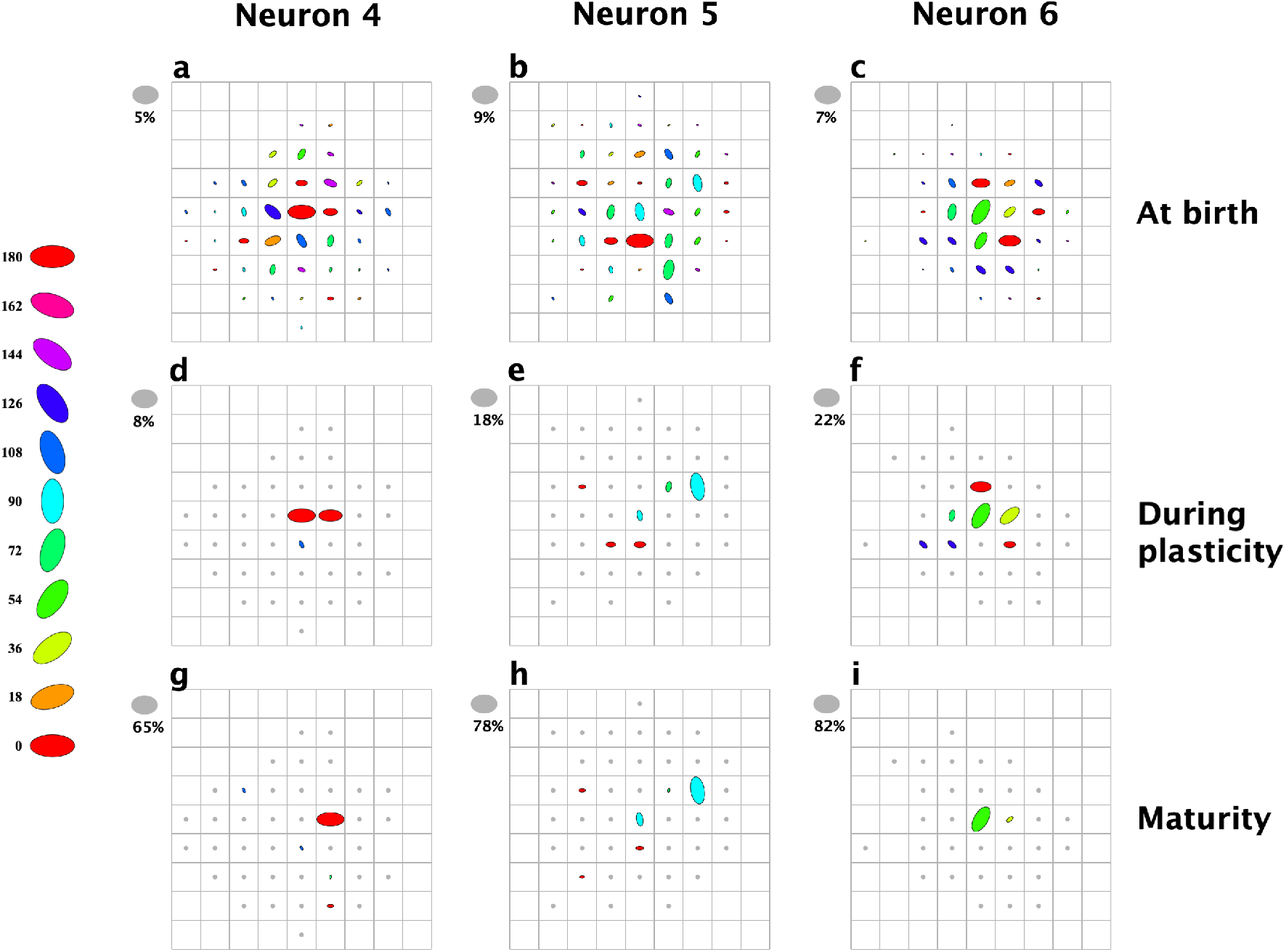
Lateral-inhibitory network: See Fig. 2 caption for further figure labelling details. The lateral-inhibitory network incorporates lateral, inhibitory synapses within V1, that are initialised to have a weight of 3, and assumed to remain constant. During development, all excitatory synapses were allowed to evolve according to the weight update equations for the lateral network, Eq. (5). Parameters governing the development of synapses between the RGC and LGN, Eq. (5a), were chosen such that all synapses reached the upper bound. Parameters for the development of synapses between the LGN and V1, Eq. (5b), were chosen such that ⅓ of the initial connections in to a V1 cell were expected to survive after the cell had reached maturity. **(a-c):** Presynaptic connections from the RGC layer to three V1 neurons, mediated by the LGN, before development. **(d-f):** The same three V1 neurons during plasticity but prior to maturity, or convergence of weights. The RGC connections to V1 are shown after 30% of the simulation has run. **(g-i):** The same three V1 neurons after convergence of weights at maturity. Figures show the RGC connections to V1 that survived development.

#### 2.2.2 Tuning of V1 cells

Tuning curves for V1 cells from the lateral-inhibitory network are shown in Figs.5a-c and 5e-g, before, during and after development, respectively. It is clear that, prior to plasticity the combined input from up to 10 orientations has been averaged to give low orientation selectivity and a corresponding preference for lower spatial frequencies. After development, the spatial frequency profile of V1 cells is markedly changed so that the peak response corresponds to the spatial frequency of the retina DOG kernels. The lateral inhibitory neurons, whose neural activity is the linear combination of many different kernel orientations covering a larger spatial extent, and hence have relatively white spatial frequency responses but overall a low spatial frequency bias, serve to suppress the response of V1 cells to all frequencies, leaving only the dominant frequency responses having any significant amplitude (i.e., band-pass, Fig. 5d). This result is markedly different to the spatial frequency responses of V1 cells from the feed-forward network, which showed a significant response to all spatial frequencies (i.e., more low-pass).

**Figure 5:**
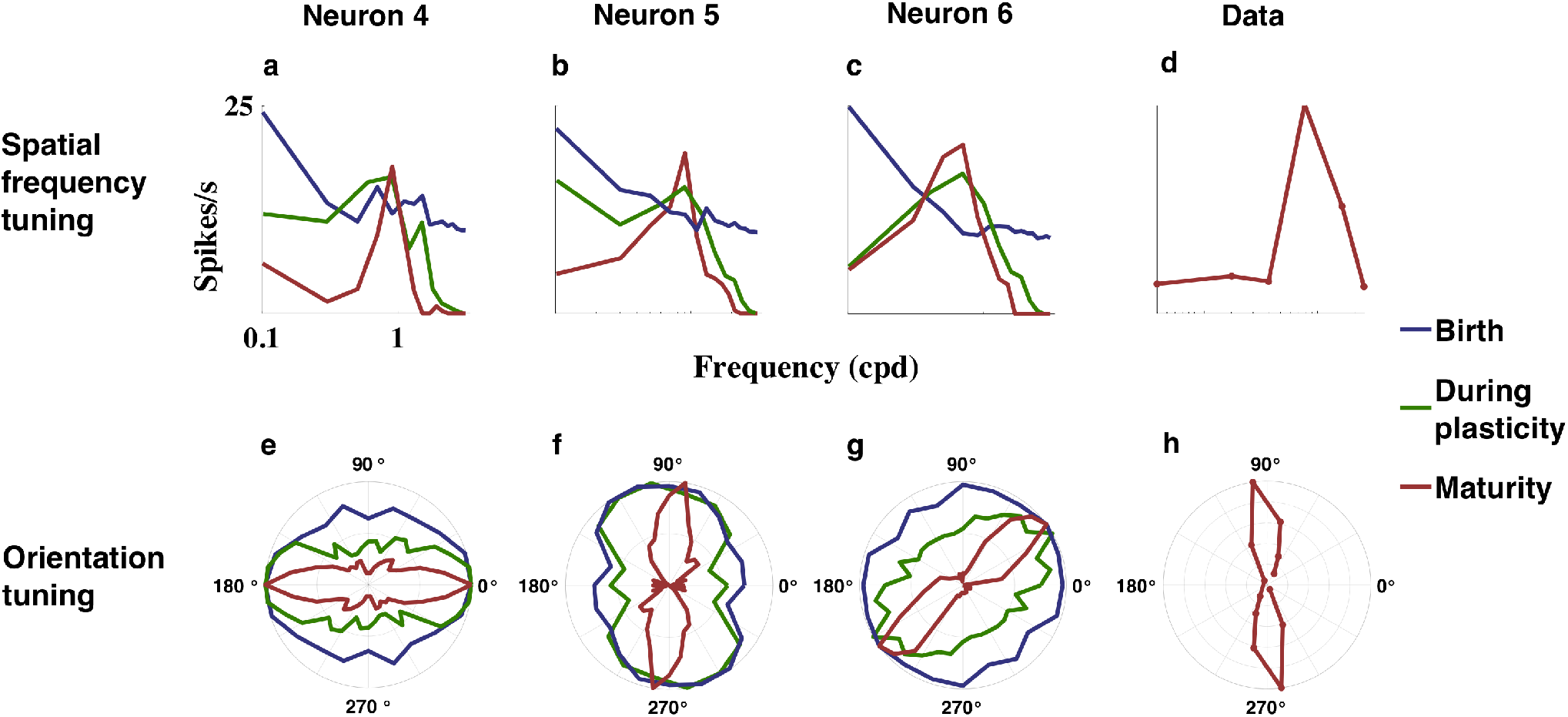
Tuning curves of V1 cells from the lateral-inhibitory network. Lateral-inhibitory network stimulated with sinusoidal gratings, to evaluate functional orientation and spatial frequency selectivities (see Fig. 3 for details). Orientation tuning curve data were normalised to 1. Spatial frequency tuning curves show a V1 cell’s firing rate as a function of the spatial frequency (in cycles per degree (CPD) of the input stimuli, at the cell’s preferred orientation. Tuning curves are shown for three exemplar cells in V1 of the lateral-inhibitory networks, prior to learning at birth (blue), during the learning process (green), and after weights have converged at maturity (red). Also included is experimental data showing the response of a cat V1 cell to a sinusoidal grating^49^. **(a-c):** Spatial frequency response of three lateral-inhibitory network V1 cells. **(d):** Spatial frequency response from a cat V1 cell^49^. **(e-g):** Normalised orientation tuning curves of three lateral-inhibitory network V1 neurons. **(h):** Normalised orientation tuning curve of a cat V1 cell.

Similar to V1 cells in the feed-forward network, after development there are only a few RGC cells inputting to a V1 cell via the LGN relay cells. A single retinal cell tended to dominate, and hence there was a preferred orientation. However, in distinction to the feed-forward network the suppression of low and high spatial frequencies renders the cell more orientation selective by reducing the CV of its orientation response. An orientation tuning curve of a real V1 cell is shown in Fig. 5h, which shows very similar CV to the model neurons.

## 3 Discussion

The phenomenon of orientation selectivity has raised two integrated but distinct questions for systems neuroscience. First, what scheme forms the neural basis of orientation selectivity, and second, how this scheme develops in the absence of visual input. Not only have both questions remained contentious, a scheme that coherently, parsimoniously and simultaneously addresses both has been lacking in the field (though see Kremkow and Alonso^24^). We propose that the critical involvement of sub-cortical biases for stimulus orientation may be such a scheme, and demonstrate computationally how this scheme of orientation selectivity can provide a solution to both the circuitry and developmental problems. Key to this demonstration was to incorporate into our model of the visual system two well established and previously overlooked aspects of the pre-cortical visual pathway: orientation biases in the sub-cortical retina and LGN, and, a dominant and divergent output in the retinogenicocortical pathway. By doing so, we have shown that orientation selectivity in primary visual cortex can come about, without any visual input driving development, through a typical plasticity process that results in V1 inheriting and amplifying the orientation selectivity from the LGN.

As for the question of what is the neural basis for orientation selectivity in mature V1, it is beyond the scope of this paper to cover an issue that has been reviewed many times^3,4,51–53^. It suffices to emphasise here the following points in support of the notion that sub-cortical biases provide an attractive answer to this question. First, the capacity of a scheme based on sub-cortical biases to generate an accurate model of V1 cells, which predicts not just orientation selectivity but other behaviours such as tuning for stimulus length and spatial frequency, has been computationally demonstrated^15^. Second, physiological studies have found congruence between LGN biases and V1 orientation selectivity^13,54^. And third, recent studies have placed a relatively low limit on the spatial extent covered by the receptive fields of LGN inputs to a single orientation domain of V1^16,55^. Such a limitation hinders the capacity of a spatial convergence scheme to generate orientation selectivity and has renewed the importance that other mechanisms, including sub-cortical biases, may have for explaining orientation selectivity^3,56,57^.

A compelling aspect of our developmental scheme is its parsimony. This arises from our scheme depending on a relatively simple pattern of correlations, and in turn, the anatomical structure behind these correlations. On the other hand, previous developmental schemes that produced orientation selectivity through spatial convergence required relatively particular patterns in the correlations between inputs to V1. Specifically, it was necessary for lateral inhibitory mechanisms to create negative correlations between visually displaced inputs of the same dark-light polarity or, positive correlations between similarly displaced inputs of opposing dark-light polarity, and for these correlations to follow a Mexican-hat curve with respect to the magnitude of the displacement^22,43,58^. Such zero-crossing Mexican-hat correlation curves were required so that once the displacement between two inputs’ receptive fields reached a certain magnitude, the correlation between them would flip their sign compared to when their source LGN cells bore receptive fields closer to each other. From this, elongated and striped input patterns emerged and imparted an orientation selective receptive field to target V1 cells.

Though Mexican-hat correlations have been reported in the mature retina^59^, this observation is not universal^60,61^. More pivotally, they are absent in the LGN during the relevant developmental period^26^. The fundamental issue with such zerocrossing correlations is likely the undue precision required of the visual circuitry at an early stage of development. The processes that would otherwise yield such correlations could be undermined by the lack of definite receptive fields^26,45^, an unfinished separation of the ON and OFF pathways^62^, and the spontaneous activity of LGN cells being influenced by endogenous and upstream sources^63^.

In contrast, the pattern of correlations required by our scheme involves only positive correlations between multiple LGN cells that share input from retina cells. In the mature visual pathway, such correlations are well established^37,64,65^, as is the necessary anatomical structure - divergent and dominant retinal inputs to LGN as in the cat, or, LGN inputs to V1 as in the macaque^28–37^. Though it is not known to what extent this can be said for the period in which orientation selectivity develops, the following points can be made. First, though the retina is not the only source of spontaneous activity in LGN, the retinal drive is likely sufficient for the targets of common retinal inputs to correlate^63,66^. And second, the sequence in which the visual pathway develops is such that orientation selectivity develops after the major functional segregations in the LGN^67^. Thus, during development of orientation selectivity, the retino-geniculate circuitry would likely be approximate to its mature state^45^ to such a degree that our required positive correlation structure is likely in place (for example, see Kiley and Usrey^45^). Indeed, as the retinogeniculate divergence is likely present early in visual development, if there are any correlations relevant to orientation selectivity present at the appropriate time, the most facile would be our required positive correlation derived from shared inputs.

A key aspect and prediction of our scheme is that each V1 cell receives inputs from only a few retinal cells. This is necessary for the sum of the inputs to bear an orientation preference. It is also the strongest difference between the sub-cortical and spatial convergence schemes. Support for this structure can be found in the substantial retina-to-V1 correlations measured in the two studies to directly observe them^38,68^ as well as the aforementioned findings of a low limit on the spatial spread of inputs to V1^16,55^. Additionally, intracellular studies have also shown that the excitatory post-synaptic potentials to single layer 4 V1 cells are restricted to a relatively small elliptical region that is not much larger than the sizes of receptive field centres of single LGN cells^12,69^. Generally, this retina-to-V1 connectivity structure is consistent with the well established functional and retinotopic precision of the inputs to V1 cells^16,55,68^. However, there is some contention concerning the degree to which a single retinal cell is responsible for the inputs to V1, with Lee et al.^38^ claiming ~ 30 – 70% and Kara and Reid^68^ ~ 3%. Apart from methodological differences in their estimates of contribution, this ~ 3% figure reported by Kara and Reid^68^, if taken alone, may underestimate an important factor in the retinal drive of V1 cells, namely temporal summation. In fact, the analysis by Kara and Reid^68^ found that two retinal spikes occurring in relatively quick succession are surprisingly efficacious at driving V1 cells. This efficacy was greater than that of individual retinal spikes and than that found previously for the combination of both the retinal drive of the LGN and the LGN drive of V1. All of this suggests that this double spike enhancement is attributable to the convergence, onto single V1 cells, of multiple LGN inputs that are synchronous due to their shared common retinal input^68,70^. Indeed, much of the contribution subtracted away by Kara and Reid^68^ in their shift prediction method, which was the key methodological difference that lead to their lower estimation of drive relative to that of Lee et al.^38^, may very well have been attributable to this double spike enhancement. The essential findings of Kara and Reid^68^ and Lee et al.^38^ may thus be more closely aligned than appears on the surface.

One outstanding issue in our scheme is the origins of the retinal orientation biases, which are thought to arise from the stretching of retinal dendritic trees as the retina matures^71^. Though it is unresolved whether these retinal biases develop by the time orientation selectivity is observed in V1, it is important to note that our developmental scheme is concerned only with anatomical connectivity and is indifferent to the receptive field structure of the retinal inputs until the time at which visual perception begins. One of its main strengths is its ability to explain the development of cortical orientation selectivity in the absence of visual experience^17,21^ from the orientation biases that are already present in the LGN prior to any visual experience^45,72^.

It may be argued that our scheme of cortical orientation selectivity being dependent on retinal biases would work perfectly with a one-to-one connectivity between retina and LGN without the need for retinogeniculate divergence that is a key component of the model proposed here. We consider that such connectivity not only takes into account the known empirical observations, but is in fact critical. RGC and LGN cell responses are nearly low pass in the spatial frequency domain and their orientation selectivity is mostly in the medium and high frequency ranges^6,73^, where the spike rate is much lower. This means that with the low spatial frequency attenuation that happens at the level of the visual cortex^49,74^, leading to the characteristic orientation and spatial frequency selectivities^73^, the remaining band-pass response needs to be boosted to yield a useful dynamic range. The divergence and convergence in the rertinogeniculate pathway is thus an essential step to enable such amplification of the spike response range in cortical cells showing orientation selectivity.

Another key difference between our scheme here and spatial convergence schemes is that we don’t consider it necessary for orientation selective V1 cells to receive monosynaptic inputs from spatially segregated LGN cells of both polarities (ON and OFF)^3^, sometimes referred to as ‘double row inputs’. This input structure is central to previous developmental schemes^58,75^, as well as to recent models of orientation selectivity^23,24^. However, that the abolition of one polarity most often doesn’t affect orientation selectivity in V1^76,77^ and that early neonatal V1 responses are orientation selective but responsive only to the OFF polarity^19^ clearly demonstrate that double row inputs are not necessary for orientation selectivity. That orientation selectivity is found in mature monopolar V1 cells^57,78^, which are probably located in the segregated polarity domains of V1^55^, adds further support to this notion. Furthermore, though developing ferrets showed some orientation selectivity without visual experience, a model based upon correlation between LGN cells could lead to sharper orientation selectivity only with visual experience^26,79^.

As Miller^22^ argues, it may be the case that double row inputs develop as an intermediate and necessary step before spatial convergence develops, such that in the mature V1 double row inputs are not necessary for orientation selectivity. However, in addition to the above comments on the issues with spatial convergence development, it is notable that our sub-cortical biases scheme also leads to a consistent and parsimonious alternative for the development of double row inputs^3^. In this alternative, specific retinal response profiles, such as for a single polarity and/or ocularity, are separated into domains across the surface of V1 with intermediate response profiles arising in the regions of V1 that are between these domains and receive overlapping inputs. This structure has been clearly demonstrated for polarity and ocularity^24^, with its geometry matching that initial predicted by our scheme^3^. Our contribution, with this scheme, is to add the orientation selectivity of retinal inputs as an additional response profile into this general process. And so, though it is beyond the scope of this paper to demonstrate the development of such V1 functional architectures, we argue that the development of double row cells, as well as binocular cells and the pinwheels of orientation selectivity, can be explained with the same parsimonious developmental scheme by which we propose basic orientation selectivity arises, probably in combination with some horizontal network mechanism^22,80,81^. Our answer, then, to Miller^22^, is that double row inputs to V1 are not universal and may develop as a side effect of our scheme.

Alternatively, spontaneous travelling retinal waves have been proposed as a source for the development of orientation selectivity ^81,82^. Such proposals, however, rely either on visual experience driving development, or at least in part^82^ (as do related sparse-coding proposals^83^), or, they rely on the development solely of the intra-cortical horizontal connections in V1, which is then postulated to later guide the plasticity of sub-cortical afferents and actual cortical receptive fields when, again, the network is driven by visual experience. Additionally, there is uncertainty around whether such waves cause activity within V1 in the appropriate fashion, if at all, and at the appropriate developmental time^22,81^. The requirement that orientation selectivity can develop without visual input^17,18,21^, as asserted earlier, is one of the central problems for all proposed developmental schemes, and one which retinal wave schemes appear not to satisfy. Our model provides, so far, the only detailed framework that can explain the development of orientation selective cells without the need for visual experience.

Our scheme has also the seeds for the development of orientation columns and the overall functional architecture of the visual cortex as described elsewhere^3^. Since specification of stimulus orientation at the retinal level is only possible, as in the case of trichromatic colour vision, with a limited number of broadly tuned channels, the cortex with its many hundreds of neurons to every retinal ganglion cell, can elaborate the whole range of preferred orientations from a limited number of co-ordinates^3,51^. This idea is supported by recent experimental work showing a possible transition from the limited co-ordinates specified sub-cortically to a range of preferred orientations at the level of the primary visual cortex^84^.

## 4 Methods

### 4.1 Network specification

To examine the impact of inhibitory connections on orientation selectivity, we consider two networks; the feed-forward network, in which all connections are excitatory, and the lateral-inhibitory network, which incorporates additional lateral inhibitory connections from the V1 inhibitory interneuron population. Cartoon figures of these networks are depicted in Fig. 1.

All excitatory connections in the network are assumed to be plastic, and bounded between 0 and 1, while connections from V1 inhibitory to V1 excitatory are assumed constant and inhibitory. While all excitatory connections are assumed plastic, connections between the retina and LGN are assumed to be unstable and therefore all converge to the upper bound (see Linsker^42^ for a discussion on model configuration required for this to occur). Similarly, connections between LGN cells and the inhibitory neurons in V1 all converge to the upper bound. Connections between the layers are assumed to be learned sequentially, so that synapses between the retina and LGN layers are learned before those between LGN and V1. This is a sensible assumption since deeper layers can only be learned after earlier layers have attained some structure.

The layers are positioned as planes lying parallel to one another, with neurons equispaced within the layer. To aid interpretation of distance metrics, we measure distances across a laminar in number of grid spaces, which is the distance between two neurons. Connections between layers have a spatial distribution such that nearby neurons in the presynaptic layer are more densely connected to a postsynaptic neuron in the next layer. As the radial distance from the neuron increases, connection density decreases as a Gaussian function of the postsynaptic distance. The two dimensional radial distance of presynaptic neuron, *u*, in the retina, from postsynaptic neuron, *n*, in layer *L*, is defined by vector, [*x_un_,y_un_*]. The density of connections from the retina to the LGN layer is parameterised by radius, (σ^RL^)^2^, which is the distance dependent variance of the Gaussian distributed connection probability, measured in grid spaces to ensure independence from cell density in the laminar. *R* denotes the retinal cell population, while L refers to the LGN layer. Consequently, the probability of presynaptic neuron, u, generating a synaptic connection to postsynaptic neuron, n, is given by

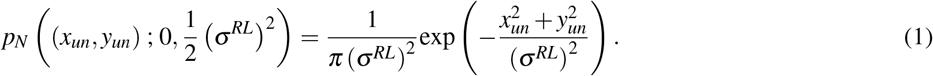

Similarly, presynaptic connections from layer *L* to layer *V*_1_ are Gaussian, parameterised by (*σ*^*LV*_1_^)^2^.

#### 4.1.1 Neuron model

The system is driven by unstructured activity from the retina, which is modelled as uncorrelated, spontaneous Poisson activity. Neural activity is assumed to be rate-based, and neurons are modelled using a Poisson neuron model. For neurons in a layer with only one neural population inputting to the layer, neural activity is updated according to the following,

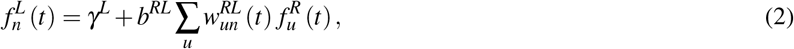

where *γ^L^* denotes the background activity of neurons in layer *L*, *b^RL^* > 0 scales presynaptic input, and 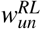 depicts synaptic strength between neurons *u* and *n*, at time *t*. Note that this model makes an implicit assumption that interlaminar distance dominates propagation delay sufficiently to assume that delay can be considered approximately equal from all presynaptic neurons to a common postsynaptic neuron.

For neurons in layer *V*_1_ that have two neural populations providing input, it’s useful to separate the sum into layer contributions, such as,

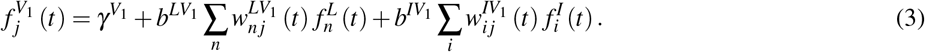

For all connections between layers, the *b* scale factor was set to the inverse of the number of presynaptic connections to each neuron. For example, for connections between *L* and *I*, *b^LI^* = 1/*N^LI^*. Set in this way, the scale factor explicitly models the limited resources that a postsynaptic neuron has to support weights and, subsequently, neural activity. Alternatively, the scaling could easily be incorporated in to the weights themselves. Also note that all neural activity was half-wave rectified so that it was always non-negative, reflecting the impossibility of a negative spike rate. Parameters were chosen such that excitatory and recurrent inhibitory inputs would balance each other out^4^.

### 4.2 Learning dynamics

Neural learning occurs very slowly, being the sum of many incremental changes. Learning occurs adiabatically slow when compared with neuronal dynamics, such as spike activity. Additionally, neurons within a neural population have the same connectivity characteristics, and the same temporal firing rate statistics. Consequently, the system can be considered ergodic. The adiabatic assumption, in conjunction with the assumption of ergodicity, allows us to write the general learning equation as a differential equation^42^. In the feed-forward network, for plastic synapses from the RGC cells, and synapses connecting the LGN and V1 layers, the learning equations are,

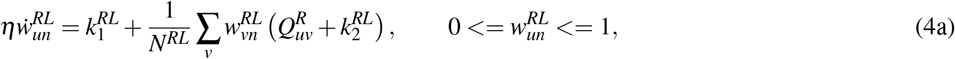

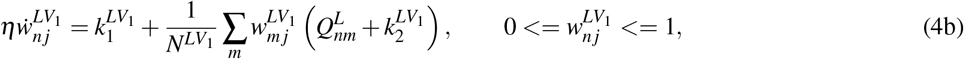

where *η* is the learning rate to ensure that learning is adiabatically slow, 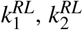, and 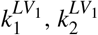, are learning constants that govern the homeostatic equilibrium, *N^RL^*, and *N*^*LV*_1_^ depict the number of synaptic connections from the relevant presynaptic layer to the postsynaptic layer, and 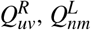 denote the normalised, expected covariance between neurons in the presynaptic layer. The origins of the learning constants, *k*_1_ and *k*_2_, are detailed in Linsker^85^, and are derived from first order statistics, including the mean spiking rate for neurons in the pre- and post-synaptic layers. They control the proportion of synapses that survive to maturity, but have no association with correlation and consequently do not influence which particular synapses survive the learning process. Inclusion of homeostatic constants is necessary to avoid all weights growing to the upper bound.

In the lateral-inhibitory network, a layer *V*_1_ neuron that receives input from two different presynaptic neural populations, *L*, and *I*, neural activity of a postsynaptic neuron is described by Eq. (3). To find the expected change in the connection strength of synapses connecting layer *L* to layer *V*_1_, we follow the method used by Linsker^42^ to get,

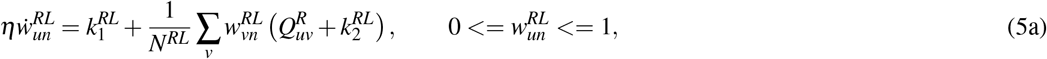

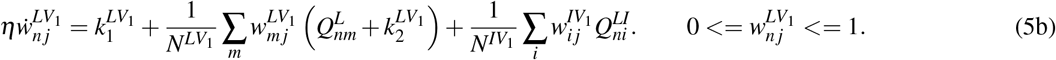

Note that this equation assumes that there are no recurrent connections between cells in the *V*_1_ layer. For excitatory synapses connecting the LGN cells to the inhibitory cells in V1, learning occurs via the Eq. (4b) learning equation, because there is only a single input layer feeding into these cells.

Linsker^42^ showed that the learning equations in Eq. (4a–5b) are unstable, and thus individual synapse weights diverge to the upper or lower bound. However, under an assumption of weak covariance in the presynaptic neural activity, stability of the mean synaptic weight depends on the values of the constants controlling homeostasis, *k*_1_ and *k*_2_ ^42^. For excitatory connections, the mean weight of synapses input to a postsynaptic cell converges to,

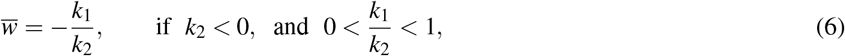

where the conditions on *k*_1_ and *k*_2_ are required to ensure that the mean synaptic weight does not diverge to the bounds.

For all synapses to grow until they reach the upper bound requires that *k*_1_ + *k*_2_ > 0. In this case the system is unstable so that the mean synaptic weight grows until all individual synapses, or all but one, have reached the upper bound^42^.

For excitatory connections between the retina and the LGN, as well as for those between the LGN and V1, we chose parameter values for *k*_1_ and *k*_2_ to ensure that the mean synaptic weight lies between 0 and 1, so that some connections grew to the upper bound, while some diminished to 0. Specifically, we chose the mean weight to be 0.25 for connections between the retina and the LGN, and 0.2 for those between the LGN and V1, which, given the upper bound of 1 on individual synapses, meant that ¼ and ⅕ of synaptic connections survived after the cell had matured, respectively. Similarly, connections between the LGN and inhibitory interneurons in the lateral network had a mean weight of 0.2 so that ¼ of synapses are expected to survive to maturity. The inhibitory connections themselves are constant and therefore do not change during learning. See Table. 1 for parameter values.

Connections to deeper layers were only allowed to evolve after connections between earlier layers had fully matured. As Linsker^42^ noted, this does not impact the final weight structure of the cells.

As addressed in the Discussion, a major distinction between our system and previous work on this problem is the pattern of correlations that drives the evolution of the system toward developing orientation selectivity. It is useful at this point to map this distinction onto the key learning equations above Eqs. (4) and (5). First, it should be noted that the learning constants *k*_1_ and *k*_2_ do not affect the structure of the correlations. As outlined above, these constants determine the average synaptic weight which, when bounded, determines the average number of, but not the structure of, synapses surviving the plasticity process. The learning constants shift the activity by the same constant value for all synapses across a layer, so as to affect the homeostasis of the system. Second, such a homeostatic constraint on the evolution of the learning equations is a necessary requirement of employing unstable learning dynamics as we do here, and has been necessary in similar work^22^. The mechanism of the constraint is not intended to emulate the corresponding biological process.

### 4.3 On-centre retina cells

On-centre cells in the retina were assumed to exhibit orientation bias, modelled using a spatial DOG kernel^86–88^. The focus on on-centre cells was motivated by parsimony, the objective being to create the simplest network that successfully explains the learning of orientation selectivity in V1. Further, combining both on- and off-centre pathways was avoided because extant schemes that rely on interactions between them, as addressed in the Introduction and Discussion, suffer from a number of issues^3,25–27,76,77^. The kernels were constant over time and, although the orientation varied across RGC cells, the extent of the bias was constant across the layer. The response of RGC cells was generated by convolving the DOG kernel with the input stimulus.

For a RGC cell, *u*, with an orientation bias of 0°, receiving an input stimulus, *S*, from location, *r*, the weight of the DOG kernel was defined as,

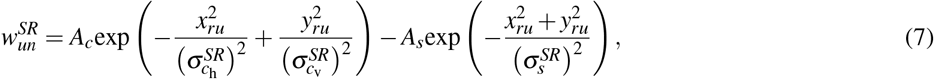

where 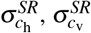, denote the standard deviations of the horizontal and vertical excitatory, centre Gaussian, while 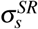, denotes the standard deviation of the symmetric, surround Gaussian. To obtain retina cells with non-zero orientation biases, we simply rotated the kernel to the desired orientation using nearest-neighbour interpolation^88^. We denote the orientation bias of a cell, *u*, in the RGC by 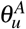. The parameters *A_c_* and *A_s_*, are scaling constants for the centre and surround Gaussians, respectively. The magnitude of the orientation biases for the above retina kernels were taken from a prior model^15^ (Fig. 6c)that was modelled on data from Vidyasagar and Urbas^8^ (Fig. 6d). The kernel with a 0° orientation bias is shown in Fig. 6b.

**Figure 6:**
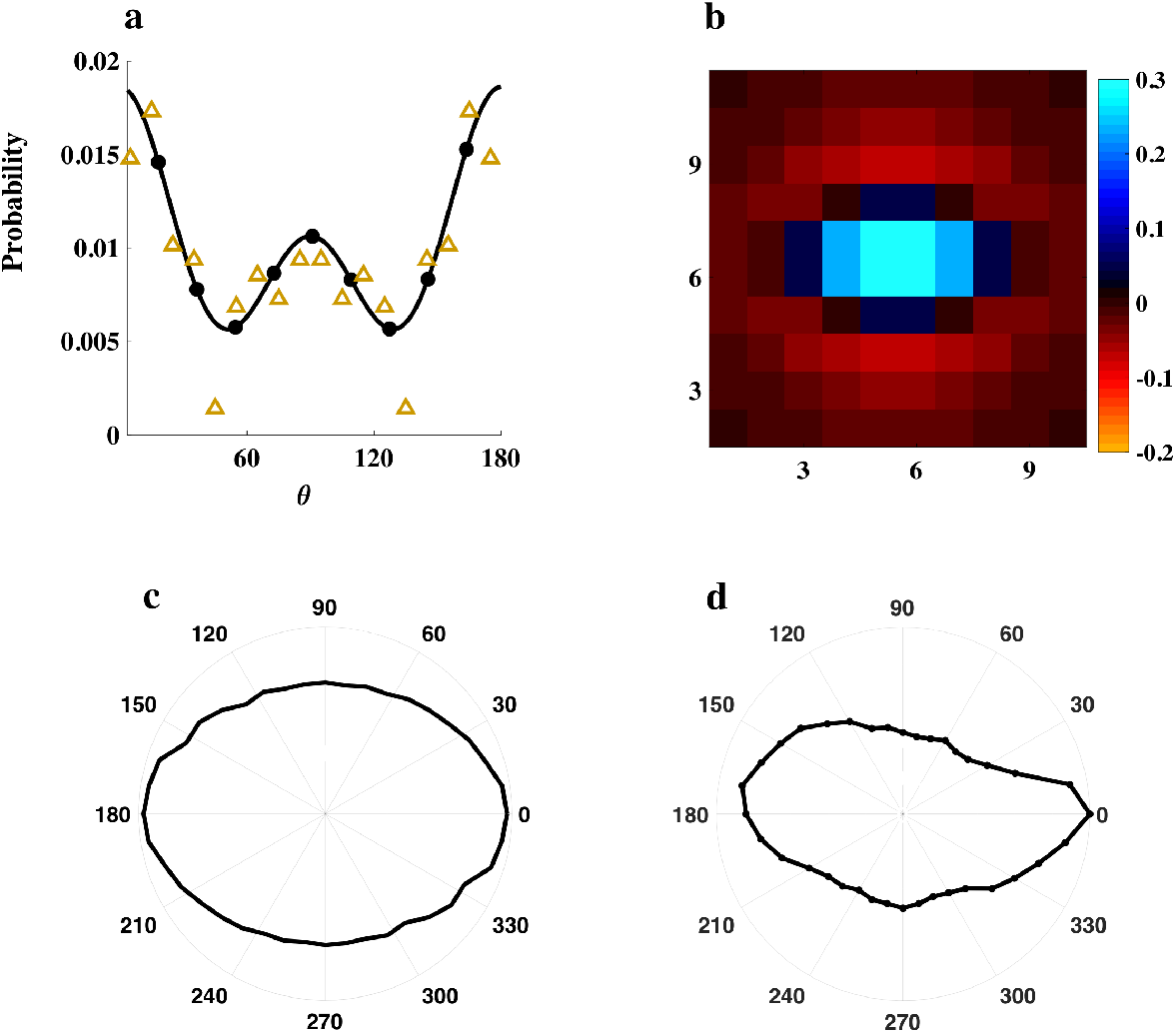
Orientation bias of the RGC cells in the retina. **(a):** Distribution of the orientation bias of RGC cells. The distribution was taken from experimental data from the cat retina^13^, resampled to extract 10 unique, equispaced orientations between 0° and 180°, shown by the dots in the figure, and rescaled to sum to 1. The orientation of a RGC cell was drawn randomly from the set of orientations according to the distribution shown. The orange triangles show the experimental data in^13^ that the distribution was generated from. **(b):** The deterministic receptive field of a RGC cell with orientation bias of 0°, constructed from DOG kernels using Eq. (7) with parameters given in Table. 1, where the horizontal and vertical axis show 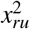 and 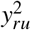, respectively. The excitatory centre is constructed from an asymmetric Gaussian, with horizontal and vertical standard deviations 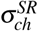, and 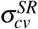, respectively, while the inhibitory surround is generated from a symmetric Gaussian with standard deviation 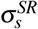 Other orientations were constructed by rotating the kernel shown using nearest-neighbour interpolation. **(c):** The orientation tuning curve of a RGC cell with an orientation bias of 0°. This figure shows the maximum firing rate of the cell for all input stimulation orientations, *θ*, using the cell’s preferred spatial frequency, *κ*, which was the frequency for which the cell responded with its highest firing rate. **(d):** Example orientation tuning curve for a real LGN cell, recorded from a cat^8^.

The distribution of orientations in the retina was modelled on data published on the dendritic orientations of cells in the cat retina, in^13^, shown in Fig. 6a. The distribution of orientations from Schall et al.^13^ was mirrored to extend to 180° and resampled to extract 10 equispaced, unique orientations between 0° and 180°, including both 90° and 180°. Given that each LGN cell received input from a single retina cell, each LGN cell had a single orientation, inherited from the retina cell.

### 4.4 Input

During the plasticity phase, the network was driven by spontaneous Poisson input from the retina, with each input retinal cell modelled as a homogeneous Poisson rate process, i.e. 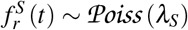, where 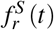 denotes the spiking rate of retinal cell *r*, and *λ_S_* is the mean spiking rate of cells in the layer over time, which is assumed constant both across the layer and over time.

Once all weights, or all but one, reached the upper or lower bound, tuning curves for cells in each layer were generated in the stimulus response phase to evaluate functional orientation and spatial frequency selectivity of cells. To this end, we stimulated the network with moving sinusoidal gratings, across a range of spatial frequencies, *κ*, and grating angles, *θ*. Input from the retinal cells to the layer LGN cells was generated according to the following,

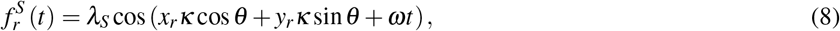

where 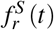 is the activity of retinal cell, *r*, at position (*x_r_*, *y_r_*) in the layer, at time *t*. The temporal frequency was constant, and set to *ω* = 2*π*. The set of grating orientations was chosen to be twice the number of input orientations, to determine if angles not available in the retina could be constructed in V1. See Table. 1 for details.

### 4.5 Evaluating Orientation selectivity (OS)

After all synapses had matured, and the plasticity phase was completed, the network was stimulated with moving sinusoidal gratings at a range of spatial frequencies, *κ* and orientations, *θ*, and a temporal frequency of 2*π*. Each grating was presented for a sufficient time that all cells in the retina experienced phases across a full period of the sinusoid.

Tuning curves for all cells in the retina and V1 were constructed by recording the cell’s maximum firing rate for every {*κ*, *θ*} combination, The spatial frequency response of a cell was determined by recording its maximum firing rate at each *κ*, using the cell’s optimal stimulation angle, *θ*. Similarly, an orientation-tuning curve for a cell was constructed by recording the cells maximal firing rate for all stimulation angles, *θ*, using the cell’s optimal spatial frequency, *κ*, after learning. Spatial frequency units were converted to physiologically realistic units of CPD of visual angle by treating the size of our RGC kernel (see Table. 1) as corresponding to the dimensions of near centre beta RGCs^28,89^.

Further, in order to highlight what receptive field changes co-occurred with the above functional changes, and that they mimicked physiological simple cell receptive fields in V1, reverse correlation receptive field mapping was simulated^90^, generating pseudo receptive fields (pseudo-RFs) for cortical cells by linearly summing the RF DOG kernels from the RGC cells, via the LGN cells, weighted by the synaptic weights between each layer, and scaled to reflect the mean firing rate of the layer (see 4.7 for example)

The preferred orientation and magnitude of selectivity of responses were quantified using conventional vector averaging of the response magnitudes for each stimulus.

### 4.6 Contrast invariance

Cortical neurons in V1 are known to exhibit contrast invariance^87,91^, in which the bias of a cell’s orientation tuning curve does not change as a function of stimulus contrast. Furthermore, a large increase in contrast elicits a much smaller change in neural response^92^. Given that contrast invariance is a key feature of V1 cells, we investigated the response of the feed-forward and lateral-inhibitory network V1 cells to stimuli with contrast increasing exponentially. The results are show in Fig. 7.

**Figure 7:**
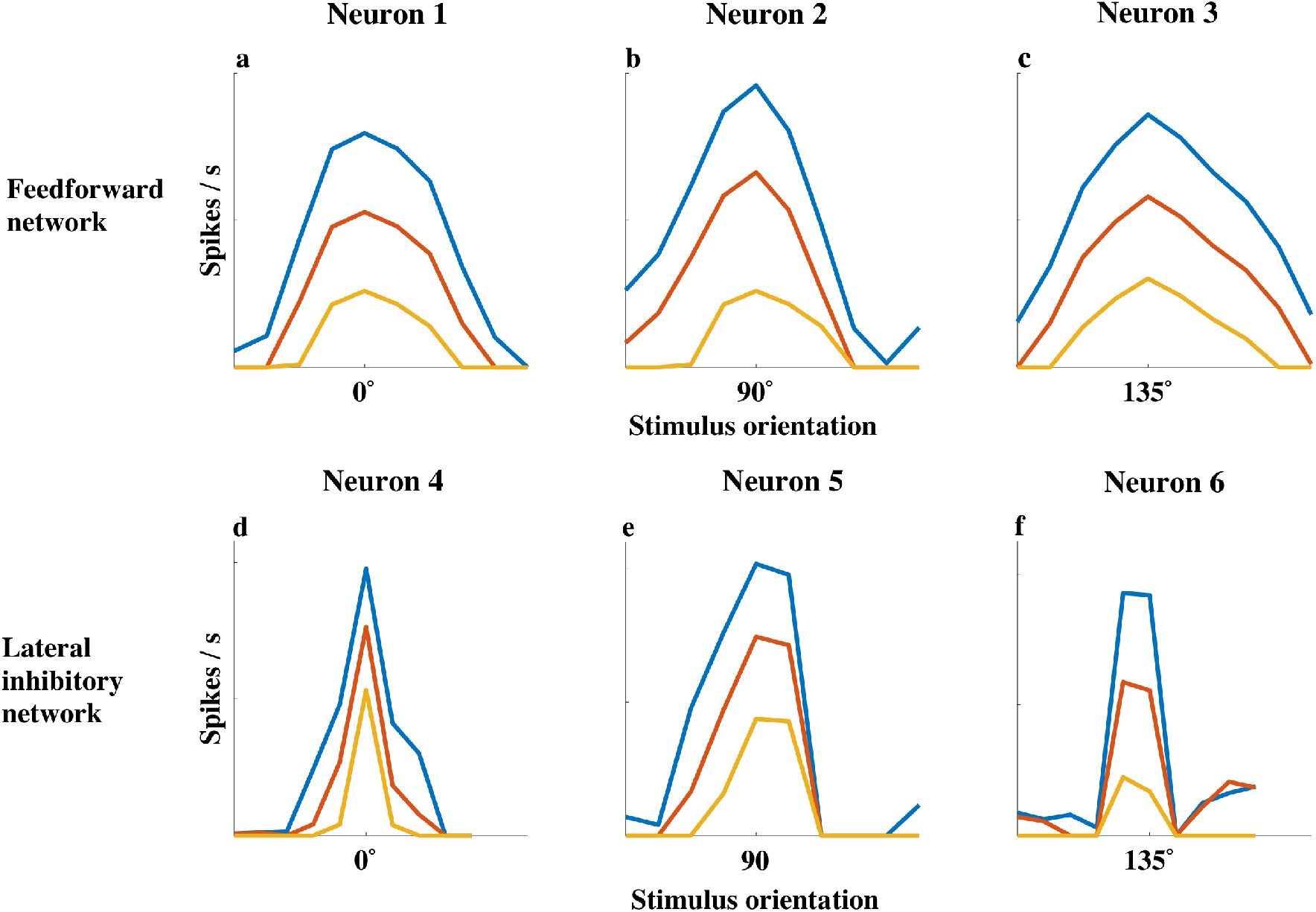
Response of V1 cells in the feed-forward and lateral-inhibitory networks to three different stimuli contrasts. Sinusoidal gratings were generated for spatial frequencies between 0 and 2.4 CPD, and for 20 orientations between 0 and 180 °. Contrast invariance was examined by generating orientation tuning curves for varying the sinusoidal amplitude, *λ_S_* in Eq. (8), between 15 (yellow), 25, and 50 (see Section 4.5 for full details). The figure depicts orientation tuning curves, generated by recording each cell’s maximum response to its preferred spatial frequency, across all input stimuli orientations. The x-axis displays the tuning curves centred around each cell’s preferred orientation, while the y-axis shows maximum firing rates in spikes/s. **(a-c):** tuning curves for cells from the feed-forward network. **(d-f):** tuning curves for cells from the lateral-inhibitory network.

Fig. 7a-c exhibits orientation tuning curves for three exemplar cells from the feed-forward network, for three different input contrasts, *λ_S_* ∈ {15,25,50}. The ‘iceberg effect’, in which the range of stimulus orientations eliciting a response in V1 cells broadens as the input contrast increases^92,93^, is clearly visible. However, as most simple cells do not respond to stimuli having an input orientation that is orthogonal to their preferred orientation, this is in contradiction to experimental findings^91,92^.

Fig. 7d-f depicts orientation tuning curves for three cells from the lateral-inhibitory network for the same three input contrasts. As the stimulus contrast increases, the range of orientations eliciting a response from V1 cells remains roughly constant, and thus the cells are considered contrast invariant^92^. The inhibitory interneurons are the mechanism behind which contrast invariance is achieved, because an increase in the activity of excitatory input is accompanied by a corresponding increasing in the activity of the interneurons. So long as the relatively white interneuron activity is sufficient to keep responses at non-optimal spatial frequencies around zero, the width of the tuning curves will remain approximately constant^94^. It should also be noted that an exponentially increasing contrast precipitates an approximately linear change in maximal firing rate, which accords with experimental findings^87,91^.

### 4.7 Pseudo receptive fields

The pseudo-RFs of V1 cells in both the feed-forward and the lateral-inhibitory networks are shown in Fig. 8. Pseudo-RFs were created by summing all of the DOG kernels from the retina that were received by a V1 cell, weighted by each of the synapses that were involved in propagating the kernel to V1. The pseudo-RFs indicate the spatial structure of the summation of retina kernels that are ultimately inputs to the cortical neurons, via the LGN cells. They therefore provide insight to the orientation tuning and spatial frequency tuning of the cortical cells and what spatial structure of inputs gives rise to these tunings, assuming that there is no temporal structure in the stimuli. These figures do not capture the impact of firing rate, and any spatial or temporal correlations in neural activity. However, they capture the structure of the kernels being input to a V1 cell. It can be seen that before plasticity much of the structure is averaged out because there are 8 to 10 orientation biases being received by the V1 cells. However, after the plasticity phase is completed, the input is dominated by a single kernel, and hence the orientation tuning is more visible.

**Figure 8:**
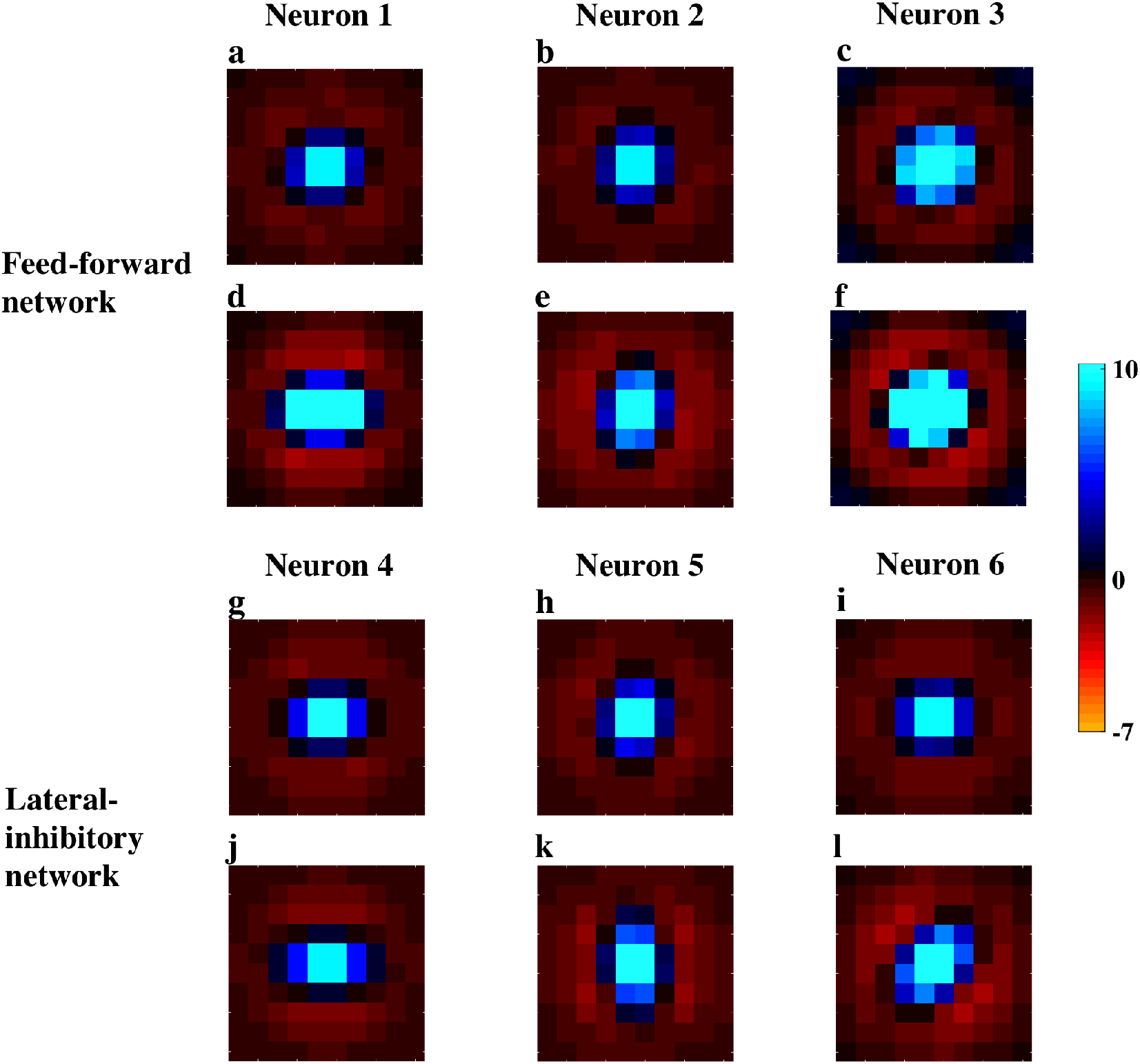
Pseudo-RFs for cortical neurons in V1, created by summing the DOG kernels from the retina, weighted by each synapse that propagates the kernel to the cell in V1. **(a,b,c):** pseudo-RFs of V1 cells in the feed-forward network prior to development. **(d,e,f):** pseudo-RFs of V1 cells in the feed-forward network after development. **(g,h,i):** pseudo-RFs of V1 cells in the lateral-inhibitory network prior to development. **(j,k,l):** pseudo-RFs of V1 cells in the lateral-inhibitory network after development.

### 4.8 Distribution of RGC inputs to V1

Fig. 9 shows a histogram of the number of unique retinal inputs to a cortical V1 cell, via the LGN, both before, during and after plasticity. The configuration of the homeostatic parameters ensures that on average only one fifth of the LGN synapses to a V1 cell survive the plasticity phase (see Eq. (6)). This is consistent for all cortical cells in both the feed-forward and lateral-inhibitory networks. Plasticity favours RGC cells that are most dominant, or have the most synaptic inputs to the V1 cell, and hence the retinal cells that have only a few connections to the V1 cells will be culled. Consequently, the histogram showing the number of unique retinal cells connected to each V1 cell demonstrates a much more significant reduction than one fifth, with the average number of unique RGC inputs to a V1 cell being 65 before plasticity, and 4 after the plasticity phase was completed.

**Figure 9:**
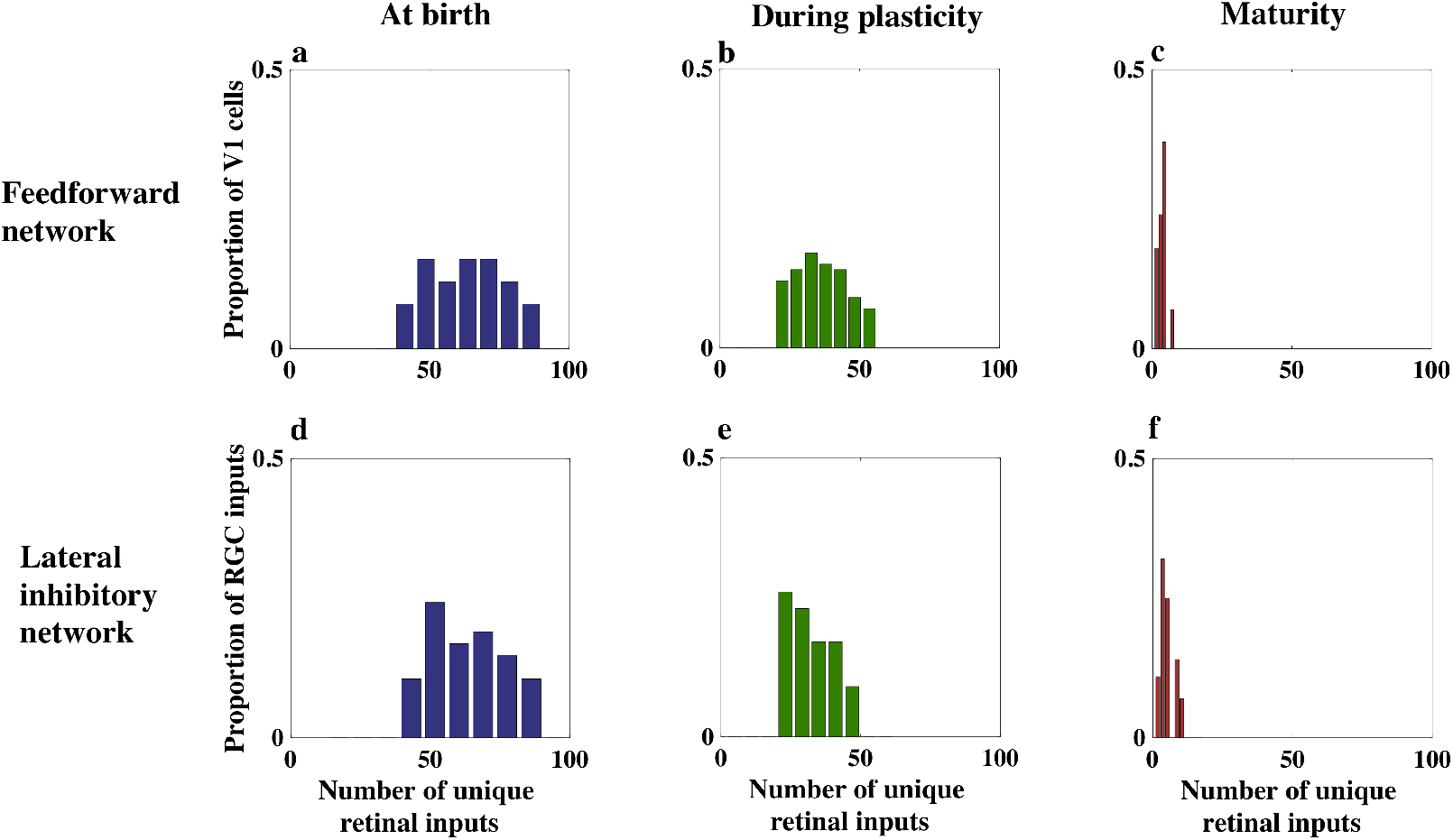
Histogram showing the number of unique retinal inputs to cortical cells in V1, via the LGN. The x-axis gives the number of unique retinal inputs, while the y-axis is the proportion of V1 cells having that number of unique retinal inputs. **(a):** At birth, for the feed-forward network. **(b):** During plasticity, for the feed-forward network. **(c):** At maturity, for the feedforward network. **(d):** At birth, for the lateral-inhibitory network. **(e):** During plasticity, for the lateral-inhibitory network. **(f):** At maturity, for the lateral-inhibitory network.

Before development there were many RGC cells with just a few synaptic connections, via the LGN, to a V1 neuron. Therefore, the proportion of total connections from each RGC cell was small. After development there was typically one RGC cell with a large proportion of the remaining synaptic connections to the cortical cell, and a few RGC cells with a smaller proportion of the synaptic connections. This tendency can be observed in Fig. 10, which shows a histogram of the proportion of synaptic connections from RGC cells to V1 cells, via the LGN.

**Figure 10:**
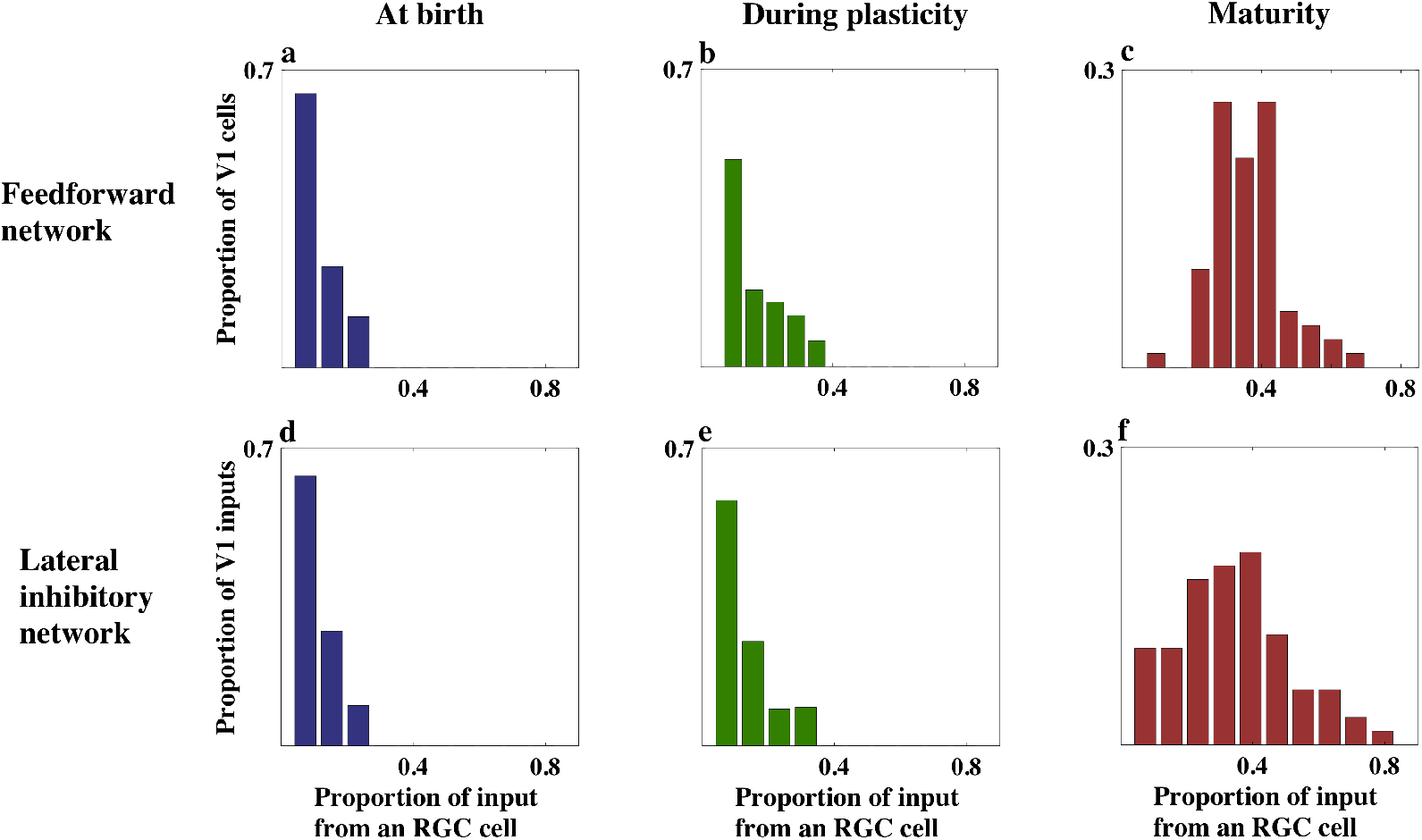
Histogram showing the proportion of total synapses a RGC cell inputs to a V1 cell via the LGN. The x-axis gives the proportion of total synapses, while the y-axis is the proportion of V1 cells having RGC inputs of that strength. **(a):** At birth, for the feed-forward network. **(b):** During plasticity, for the feed-forward network. **(c):** At maturity, for the feed-forward network. **(d):** At birth, for the lateral-inhibitory network. **(e):** During plasticity, for the lateral-inhibitory network. **(f):** At maturity, for the lateral-inhibitory network.

## 5 Data Availability

Part of the data generated or analysed during this study are included in the published article. The rest are available from the corresponding author or Catherine E. Davey on reasonable request.

## Acknowledgement

Support under the Australian Research Councils Discovery Projects funding scheme is acknowledged by CED and ANB (DP140102947) and TRV (DP DP170104170).

## Author contributions

TRV and EKJL conceptualised the basic model, CED, EKJL and LK designed the model, CED wrote the code for simulation, all authors refined the models and designed experiments, CED, EKJL and TRV drafted the paper, which all authors edited and gave feedback, and ANB and TRV obtained funding.

## References

[1] Hubel, D. H. & Wiesel, T. N. Receptive fields, binocular interaction and functional architecture in the cat’s visual cortex. The Journal of Physiology 160, 106–154 (1962).

[2] Hubel, D. H. & Wiesel, T. N. Receptive fields and functional architecture of monkey striate cortex. The Journal of Physiology 195, 215–243 (1968).

[3] Vidyasagar, T. R. & Eysel, U. T. Origins of feature selectivities and maps in the mammalian primary visual cortex. Trends in neurosciences 38, 475–485 (2015).

[4] Priebe, N. J. Mechanisms of Orientation Selectivity in the Primary Visual Cortex. Annual Review of Vision Science 2, 85–107 (2016).

[5] Sedigh-Sarvestani, M. et al. Intracellular, In Vivo, Dynamics of Thalamocortical Synapses in Visual Cortex. Journal of Neuroscience 37, 5250–5262 (2017).

[6] Levick, W. R. & Thibos, L. N. Orientation bias of cat retinal ganglion cells. Nature 286, 389–390 (1980).

[7] Passaglia, C. L. & Lee, B. B. Orientation sensitivity of ganglion cells in primate retina. Vision Research 42, 683–694 (2002).

[8] Vidyasagar, T. R. & Urbas, J. V. Orientation sensitivity of cat LGN neurones with and without inputs from visual cortical areas 17 and 18. Experimental Brain Research 46, 157–169 (1982).

[9] Shou, T. D. & Leventhal, A. G. Organized arrangement of orientation-sensitive relay cells in the cat’s dorsal lateral geniculate nucleus. The Journal of Neuroscience 9, 4287–4302 (1989).

[10] Vidyasagar, T. R., Pei, X. & Volgushev, M. Multiple mechanisms underlying the orientation selectivity of visual cortical neurones. Trends in neurosciences 19, 272–277 (1996).

[11] Vidyasagar, T.R. Subcortical mechanisms in orientation sensitivity of cat visual cortical cells. Neuroreport 3, 185–188 (1992).

[12] Pei, X., Vidyasagar, T.R., Volgushev, M. & Creutzfeldt, O. D. Receptive field analysis and orientation selectivity of postsynaptic potentials of simple cells in cat visual cortex. Journal of Neuroscience 14, 7130–7140 (1994).

[13] Schall, J. D., Vitek, D. J. & Leventhal, A. G. Retinal constraints on orientation specificity in cat visual cortex. The Journal of Neuroscence 6, 823–836 (1986).

[14] Viswanathan, S., Jayakumar, J. & Vidyasagar, T. R. Role of feedforward geniculate inputs in the generation of orientation selectivity in the cat’s primary visual cortex. The Journal of Physiology 589, 2349–2361 (2011).

[15] Kuhlmann, L. & Vidyasagar, T. R. A computational study of how orientation bias in the lateral geniculate nucleus can give rise to orientation selectivity in primary visual cortex. Frontiers in Systems Neuroscience 5, 81 (2011).

[16] Jin, J., Wang, Y., Swadlow, H. A. & Alonso, J. M. Population receptive fields of ON and OFF thalamic inputs to an orientation column in visual cortex. Nature Neuroscience 14, 232–238 (2011).

[17] Hubel, D. H. & Wiesel, T. N. Receptive Fields Of Cells In Striate Cortex Of Very Young, Visually Inexperienced Kittens. Journal of Neurophysiology 26, 994–1002 (1963).

[18] Wiesel, T. N. & Hubel, D. H. Ordered arrangement of orientation columns in monkeys lacking visual experience. Journal of comparative neurology 158, 307–318 (1974).

[19] Albus, K. & Wolf, W. Early post-natal development of neuronal function in the kitten’s visual cortex: a laminar analysis. Journal of Physiology 348, 153–185 (1984).

[20] Braastad, B. O. & Heggelund, P. Development of spatial receptive-field organization and orientation selectivity in kitten striate cortex. Journal of Neurophysiology 53, 1158–1178 (1985).

[21] Crair, M. C., Gillespie, D. C. & Stryker, M. P. The role of visual experience in the development of columns in cat visual cortex. Science (New York, N.Y.) 279, 566–570 (1998).

[22] Miller, K. D. A model for the development of simple cell receptive fields and the ordered arrangement of orientation columns through activity-dependent competition between ON-and OFF-center inputs. The Journal of Neuroscience 14, 409–441 (1994).

[23] Paik, S.-B. & Ringach, D. L. Retinal origin of orientation maps in visual cortex. Nature Publishing Group 14, 919–925 (2011).

[24] Kremkow, J. & Alonso, J.-M. Thalamocortical Circuits and Functional Architecture. Annual Review of Vision Science (2018).

[25] Martin, K. A. & Whitteridge, D. Form, function and intracortical projections of spiny neurones in the striate visual cortex of the cat. The Journal of Physiology (1984).

[26] Ohshiro, T. & Weliky, M. Simple fall-off pattern of correlated neural activity in the developing lateral geniculate nucleus. Nature Neuroscience 9, 1541–1548 (2006).

[27] Schottdorf, M., Keil, W., Coppola, D., White, L. E. & Wolf, F. Random Wiring, Ganglion Cell Mosaics, and the Functional Architecture of the Visual Cortex. PLoS Computational Biology 11, e1004602 (2015).

[28] Peichl, L. & Wässle, H. Size, scatter and coverage of ganglion cell receptive field centres in the cat retina. The Journal of Physiology 291, 117 (1979).

[29] Peters, A. & Payne, B. R. Numerical Relationships between Geniculocortical Afferents and Pyramidal Cell Modules in Cat Primary Visual Cortex. Cerebral Cortex 3, 69–78 (1993).

[30] Peters, A., Payne, B. R. & Budd, J. A numerical analysis of the geniculocortical input to striate cortex in the monkey. Cerebral Cortex 4, 215–229 (1994).

[31] Cleland, B. G. & Lee, B. B. A comparison of visual responses of cat lateral geniculate nucleus neurones with those of ganglion cells afferent to them. Journal of Physiology 369, 249–268 (1985).

[32] Usrey, W. M., Reppas, J. B. & Reid, R. C. Paired-spike interactions and synaptic efficacy of retinal inputs to the thalamus. Nature 395, 384–387 (1998).

[33] Usrey, W. M. & Reid, R. C. Synchronous activity in the visual system. Annual review of physiology 61, 435–456 (1999).

[34] Sincich, L. C., Adams, D. L., Economides, J. R. & Horton, J. C. Transmission of spike trains at the retinogeniculate synapse. Journal of Neuroscience 27, 2683–2692 (2007).

[35] Friedlander, M. J., Lin, C. S., Stanford, L. R. & Sherman, S. M. Morphology of functionally identified neurons in lateral geniculate nucleus of the cat. Journal of Neurophysiology 46, 80–129 (1981).

[36] Hamos, J. E., Van Horn, S. C., Raczkowski, D. & Sherman, S. M. Synaptic circuits involving an individual retinogeniculate axon in the cat. Journal of Comparative Neurology 259, 165–192 (1987).

[37] Yeh, C.-I., Stoelzel, C. R., Weng, C. & Alonso, J. M. Functional Consequences of Neuronal Divergence Within the Retinogeniculate Pathway. Journal of Neurophysiology 101, 2166–2185 (2009).

[38] Lee, B. B., Cleland, B. G. & Creutzfeldt, O. D. The retinal input to cells in area 17 of the cat’s cortex. Experimental Brain Research 30 (1977).

[39] Schall, J. D., Perry, V. H. & Leventhal, A. G. Retinal ganglion cell dendritic fields in old-world monkeys are oriented radially. Brain research 368, 18–23 (1986).

[40] Alonso, J. M., Yeh, C.-I., Weng, C. & Stoelzel, C. Retinogeniculate connections: A balancing act between connection specificity and receptive field diversity. Progress in Brain Research 154, 3–13 (2006).

[41] Dacey, D. M., Peterson, B. B., Robinson, F. R. & Gamlin, P. D. Fireworks in the primate retina: in vitro photodynamics reveals diverse LGN-projecting ganglion cell types. Neuron 37, 15–27 (2003).

[42] Linsker, R. From basic network principles to neural architecture: emergence of spatial-opponent cells. PNAS 83, 7508–7512 (1986).

[43] Linsker, R. From basic network principles to neural architecture: emergence of orientation-selective cells. PNAS 83, 8390–8394 (1986).

[44] Linsker, R. From basic network principles to neural architecture: emergence of orientation columns. PNAS 83, 8779–8783 (1986).

[45] Kiley, C. W. & Usrey, W. M. Orientation Tuning of Correlated Activity in the Developing Lateral Geniculate Nucleus. Journal of Neuroscience 37, 11549–11558 (2017).

[46] Berardi, N. & Morrone, M. C. Development of gamma-aminobutyric acid mediated inhibition of x cells of the cat lateral geniculate nucleus. The Journal of Physiology 357, 525–537 (1984).

[47] Daniels, J. D., Pettigrew, J. D. & Norman, J. L. Development of single-neuron responses in kitten’s lateral geniculate nucleus. Journal of Neurophysiology 41, 1373–1393 (1978).

[48] Wolf, W., Hicks, T. & Albus, K. The contribution of GABA-mediated inhibitory mechanisms to visual response properties of neurons in the kitten’s striate cortex. Journal of Neuroscience 6, 2779–2795 (1986).

[49] Vidyasagar, T. R. & Mueller, A. Function of GABAA inhibition in specifying spatial frequency and orientation selectivities in cat striate cortex. Experimental Brain Research 98, 31–38 (1994).

[50] Vidyasagar, T. R. Contribution of inhibitory mechanisms to the orientation sensitivity of cat dLGN Neurones. Experimental Brain Research 55, 192–195 (1984).

[51] Vidyasagar, T. R. A model of striate response properties based on geniculate anisotropies. Biological cybernetics 57, 11–23 (1987).

[52] Sompolinsky, H. & Shapley, R. New perspectives on the mechanisms for orientation selectivity. Current Opinion in Neurobiology 7, 514–522 (1997).

[53] Ferster, D. & Miller, K. D. Neural mechanisms of orientation selectivity in the visual cortex. Neuroscience 23, 441–471 (2000).

[54] Vidyasagar, T. R., Jayakumar, J., Lloyd, E. & Levichkina, E. V. Subcortical orientation biases explain orientation selectivity of visual cortical cells. Physiological reports 3 (2015).

[55] Jin, J. Z. et al. On and off domains of geniculate afferents in cat primary visual cortex. Nature Neuroscience 11, 88–94 (2008).

[56] Stanley, G. B. et al. Visual orientation and directional selectivity through thalamic synchrony. Journal of Neuroscience 32, 9073–9088 (2012).

[57] Kremkow, J., Jin, J., Wang, Y. & Alonso, J. M. Principles underlying sensory map topography in primary visual cortex. Nature 533, 52–57 (2016).

[58] Miller, K. D., Erwin, E. & Kayser, A. Is the development of orientation selectivity instructed by activity? Journal of Neurobiology (1999).

[59] Meister, M., Lagnado, L. & Baylor, D. A. Concerted signaling by retinal ganglion cells. Science 270, 1207–1210 (1995).

[60] Greschner, M. et al. Correlated firing among major ganglion cell types in primate retina. The Journal of Physiology 589, 75–86 (2011).

[61] Shlens, J. et al. The structure of multi-neuron firing patterns in primate retina. Journal of Neuroscience 26, 8254–8266 (2006).

[62] Bodnarenko, S. R., Jeyarasasingam, G. & Chalupa, L. M. Development and regulation of dendritic stratification in retinal ganglion cells by glutamate-mediated afferent activity. The Journal of Neuroscience 15, 7037–7045 (1995).

[63] Weliky, M. & Katz, L. C. Correlational structure of spontaneous neuronal activity in the developing lateral geniculate nucleus in vivo. Science (New York, N.Y.) 285, 599–604 (1999).

[64] Arnett, D. W. Correlation analysis of units recorded in the cat dorsal lateral geniculate nucleus. Experimental Brain Research 24, 111–130 (1975).

[65] Alonso, J. M., Usrey, W. M. & Reid, R. C. Precisely correlated firing in cells of the lateral geniculate nucleus. Nature 383, 815–819 (1996).

[66] Mooney, R., Penn, A. A., Gallego, R. & Shatz, C. J. Thalamic relay of spontaneous retinal activity prior to vision. Neuron 17, 863–874 (1996).

[67] Huberman, A. D., Feller, M. B. & Chapman, B. Mechanisms Underlying Development of Visual Maps and Receptive Fields. Annual review of neuroscience 31, 479–509 (2008).

[68] Kara, P. & Reid, R. C. Efficacy of retinal spikes in driving cortical responses. Journal of Neuroscience 23, 8547–8557 (2003).

[69] Creutzfeldt, O. D., Kuhnt, U. & Benevento, L. A. An intracellular analysis of visual cortical neurones to moving stimuli: Responses in a co-operative neuronal network. Experimental Brain Research 21, 251–274 (1974).

[70] Reid, R. C. Divergence and reconvergence: multielectrode analysis of feedforward connections in the visual system. Progress in Brain Research 130, 141–154 (2001).

[71] Schall, J. D. & Leventhal, A. G. Relationships between ganglion cell dendritic structure and retinal topography in the cat. Journal of Comparative Neurology 257, 149–159 (1987).

[72] Albus, K., Wolf, W. & Beckmann, R. Orientation bias in the response of kitten LGNd neurons to moving light bars. Brain research 282, 308–313 (1983).

[73] Vidyasagar, T. R. & Heide, W. Geniculate orientation biases seen with moving sine wave gratings: implications for a model of simple cell afferent connectivity. Experimental Brain Research 57, 196–200 (1984).

[74] Movshon, J. A., Thompson, I. D. & Tolhurst, D. J. Spatial and temporal contrast sensitivity of neurones in areas 17 and 18 of the cat’s visual cortex. Journal of Physiology 283, 101–120 (1978).

[75] Chapman, B. & Gödecke, I. Cortical cell orientation selectivity fails to develop in the absence of ON-center retinal ganglion cell activity. Journal of Neuroscience 20, 1922–1930 (2000).

[76] Sherk, H. & Horton, J. C. Receptive field properties in the cat’s area 17 in the absence of on-center geniculate input. The Journal of Neuroscience 4, 381–393 (1984).

[77] Schiller, P. H. Central connections of the retinal ON and OFF pathways. Nature 297, 580–583 (1982).

[78] Bishop, P. O., Coombs, J. S. & Henry, G. H. Responses to visual contours: spatio-temporal aspects of excitation in the receptive fields of simple striate neurones. The Journal of Physiology 219, 625–657 (1971).

[79] Ohshiro, T., Hussain, S. & Weliky, M. Development of cortical orientation selectivity in the absence of visual experience with contour. Journal of Neurophysiology 106, 1923–1932 (2011).

[80] Kaschube, M. et al. Universality in the evolution of orientation columns in the visual cortex. Science (New York, N.Y.) 330, 1113–1116(2010).

[81] Grabska-Barwińska, A. & von der Malsburg, C. Establishment of a scaffold for orientation maps in primary visual cortex of higher mammals. Journal of Neuroscience 28, 249–257 (2008).

[82] Hunt, J. J., Ibbotson, M. & Goodhill, G. J. Sparse coding on the spot: Spontaneous retinal waves suffice for orientation selectivity. Neural computation 24, 2422–2433 (2012).

[83] Olshausen, B. A. & Field, D. J. Emergence of simple-cell receptive field properties by learning a sparse code for natural images. Nature 381, 607–609 (1996).

[84] Mohan, Y. S., Jayakumar, J., Lloyd, E. K., Levichkina, E. & Vidyasagar, T. R. Diversity of feature selectivity in macaque visual cortex arising from a limited number of broadly tuned input channels. Cerebral Cortex 29, 5255–5268 (2019).

[85] Linsker, R. From basic network principles to neural architecture: Emergence of spatial-opponent cells. Neurobiology 83, 7508–7512 (1986).

[86] Soodak, R. E. The retinal ganglion cell mosaic defines orientation columns in striate cortex. PNAS 84, 3936–3940 (1987).

[87] Troyer, T. W., Krukowski, A. E., Priebe, N. J. & Miller, K. D. Contrast-Invariant Orientation Tuning in Cat Visual Cortex: Thalamocortical Input Tuning and Correlation-Based Intracortical Connectivity. Journal of Neuroscience 18, 5908–5927 (1998).

[88] Teich, A. F. & Qian, N. Learning and adaptation in a recurrent model of V1 orientation selectivity. Journal of Neurophysiology 89, 2086–2100 (2003).

[89] Wässle, H., Boycott, B. B. & Illing, R. B. Morphology and Mosaic of on-and off-Beta Cells in the Cat Retina and Some Functional Considerations. Proceedings of the Royal Society of London. Series B, Biological Sciences 212, 177–195 (1981).

[90] Ringach, D. & Shapley, R. Reverse correlation in neurophysiology. Cognitive Science 28, 147–166 (2004).

[91] Finn, I. M., Priebe, N. J. & Ferster, D. D. The Emergence of Contrast-Invariant Orientation Tuning in Simple Cells of Cat Visual Cortex. Neuron 54, 16–16 (2007).

[92] Carandini, M. Melting the iceberg: contrast invariance in visual cortex. Neuron 54, 11–13 (2007).

[93] Wehr, M. & Zador, A. M. Balanced inhibition underlies tuning and sharpens spike timing in auditory cortex. Nature Publishing Group 426, 442–446 (2003).

[94] Katzner, S., Busse, L. & Carandini, M. GABAA inhibition controls response gain in visual cortex. The Journal of Neuroscience 31, 5931–5941 (2011).

[95] Teich, A. F. & Qian, N. Comparison among some models of orientation selectivity. Journal of Neurophysiology 96, 404–419 (2006).

[96] Soodak, R. E., Shapley, R. M. & Kaplan, E. Linear mechanism of orientation tuning in the retina and lateral geniculate nucleus of the cat. Journal of Neurophysiology 58, 267–275 (1987).

[97] Soodak, R. E. Two-dimensional modeling of visual receptive fields using Gaussian subunits. PNAS 83, 9259–9263 (1986).

[98] Antonini, A. & Stryker, M. P. Antonini, Stryker-Development of individual geniculocortical arbors in cat striate cortex and effects of binocular impulse blockade.-1993-J Neurosci. The Journal of Neuroscience 13, 3549–3573 (1990).

[99] Yousef, T. T. et al. Orientation topography of layer 4 lateral networks revealed by optical imaging in cat visual cortex (area 18). European Journal of Neuroscience 11, 4291–4308 (1999).

[100] Crook, J. M., Kisvárday, Z. F. & Eysel, U. T. Evidence for a contribution of lateral inhibition to orientation tuning and direction selectivity in cat visual cortex: reversible inactivation of functionally characterized sites combined with neuroanatomical tracing techniques. European Journal of Neuroscience 10, 2056–2075 (1998).

